# 3D MERFISH reveals cell-type diversity generated by mRNA partitioning in spiralian embryogenesis

**DOI:** 10.64898/2026.10.05.756875

**Authors:** Yiqun Wang, Tejasvi Patil, Stephanie Neal, Ignatius Jenie, Emily Ignatoff, Bogdan Bintu, Deirdre C. Lyons

## Abstract

Cell type diversification during development is often attributed to differential gene transcription. Here, we show that, during *Crepidula atrasolea* embryogenesis, transcriptomically distinct cell types arise through differential distribution of pre-existing mRNAs. Combining immunofluorescence, live imaging, and whole-mount 3D spatial transcriptomics (MERFISH) of over 40 embryos, we show that many mRNAs form subcellular aggregates during mitosis, which segregate asymmetrically between daughter cells through microtubule associations. Cell divisions thus simultaneously increase cell number and diversity, producing six transcriptomically distinct cell quartets in the 24-cell stage embryo. We modeled aggregate compositions across cell divisions using transcript abundance and quantified transcript-specific affinity. These results establish mRNA sorting and inheritance as a scalable cell type patterning mechanism and provide a framework for investigating how mRNA segregation shapes animal development.

## Introduction

During metazoan embryogenesis, a single zygotic cell is transformed into diverse cell types with specialized morphologies, functions, and molecular identities. Single-cell transcriptomic techniques have facilitated the characterization of embryonic cell types in numerous organisms, charting the progressive emergence of distinct transcriptomic states and linking them to cell fate and function (*1–7*). These studies have established when transcriptomic differences arise among embryonic cells, but the mechanisms that generate these differences remain incompletely understood.

Differences in cells’ transcriptomic profiles are largely attributed to transcriptional regulation (*8, 9*). In this framework, genes are transcribed at different levels in different cells. However, cells can also acquire different transcriptomic profiles through the unequal inheritance of pre-existing mRNAs, as was demonstrated for a subset of genes across different species. For example, in fish and frog zygotes, germ cell determinant mRNAs, such as *vasa* and *nanos*, are restricted to a small cytoplasmic compartment. Subsequent cell divisions progressively segregate this compartment into a small subset of cells, generating a cell population enriched with germline determinants (*10, 11*). In the early mouse embryo, asymmetric cortical association of *cdx2* mRNAs results in its differential inheritance between the inner and outer daughter cells (*12*). These and other examples demonstrate that subcellularly localized transcripts can be asymmetrically inherited during embryogenesis and influence developmental patterning (*13–18*). However, developmental studies have generally examined the localization of a handful of transcripts one at a time. Therefore, it remains unclear whether asymmetric inheritance may act broadly on larger populations of transcripts to generate differences among embryonic lineages and, if so, how distinct populations of mRNAs are selected for asymmetric inheritance across lineages. Addressing these questions requires both a suitable model system and methods capable of simultaneously measuring transcript identity, subcellular localization, and cell lineage context across large numbers of genes.

Spiralian embryogenesis is an excellent model system for investigating cell type diversification with tractable cell lineages. Spiralian embryos follow a highly stereotyped cleavage program, termed “spiral cleavage”, in which cells with defined lineages and fates arise in reproducible sequences and positions (*19–21*). Spiralians are the largest of the three major bilaterian clades, comprising over half of bilaterian phyla, including molluscs, annelids, nemerteans, and flatworms (**Fig. 1A**) (*21–23*). Once foundational to experimental embryology, spiralians remain poorly represented among modern molecular model systems, leaving the mechanisms underlying spiral cleavage comparatively unexplored. Because spiral cleavage is thought to be ancestral to this entire clade (*24*), and modifications to this program are thought to contribute substantially to spiralian body-plan diversity (*21*); understanding its mechanistic basis stands to illuminate a large, ancient, and underexplored way of being an animal.

**Fig. 1.**
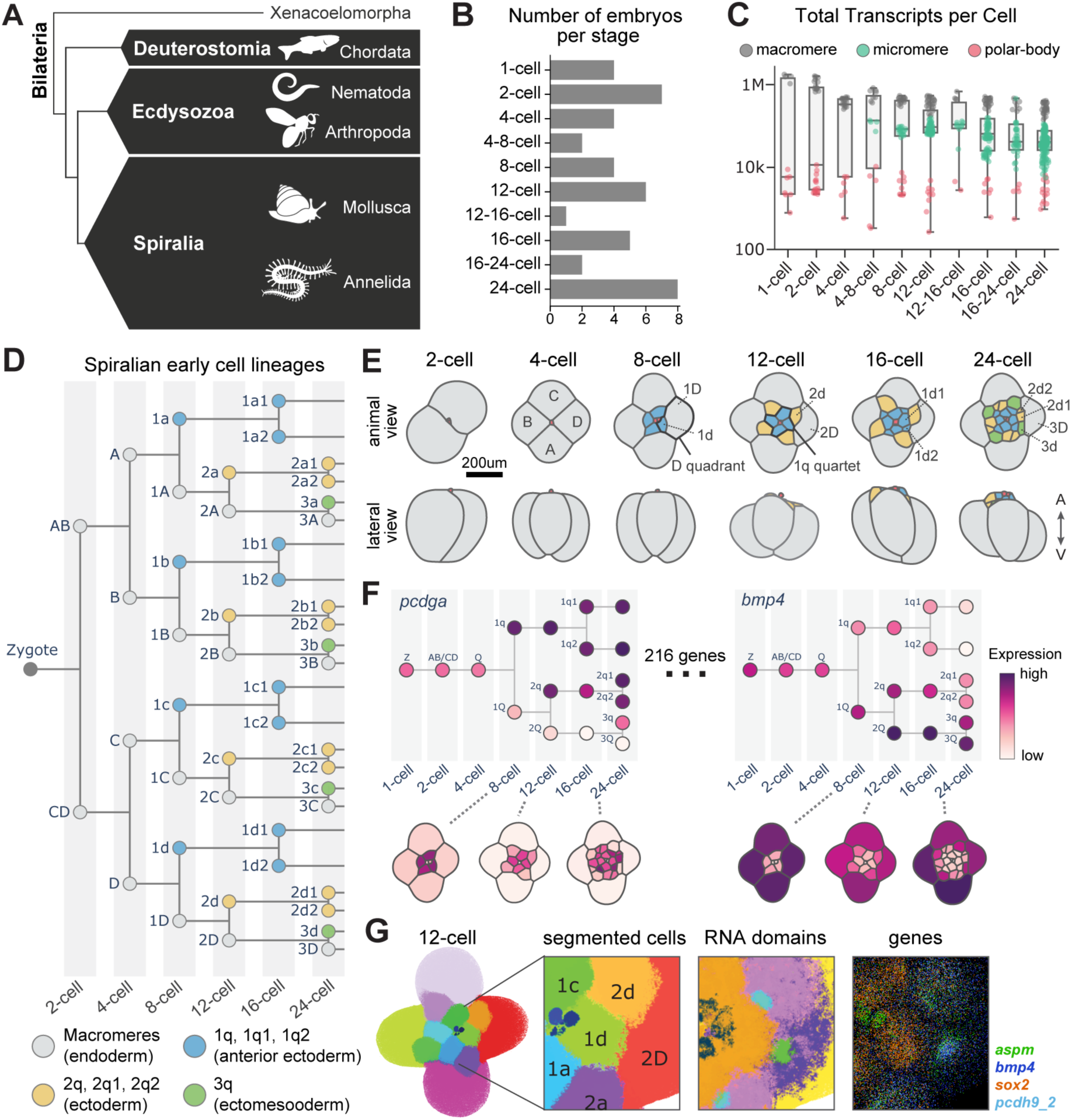
MERFISH on whole-mount embryos across early cleavage stages resolves gene expression across lineage, cell, and subcellular domains. (**A**) Phylogenetic tree showing the major clades in Bilateria. Example phyla are shown for each clade. The height of boxes are proportional to the number of phyla in each clade. (**B**) Number of embryos assayed by MERFISH across developmental stages. (**C**) Number of total transcripts passed quality check per cell. Boxes show the interquartile range (25th-75th percentile) with solid lines marking the median. Whiskers extend to the most extreme data points within 1.5x the interquartile range. Points show individual cells. Polar bodies are small non-embryonic cells formed during oogenesis. (**D**) Cell lineages of spiral cleaving embryos up to the 24-cell stage. Nodes are cells and are annotated with standard nomenclature. Colors separate major lineages. (**E**) Illustrations of representative embryos from the MERFISH assay with cell lineages annotated. Cells are colored by their major lineages as in **D**. “A” - animal pole; “V” - vegetal pole. (**F**) Cell segmentation and annotation allowed mapping of MERFISH expression to both the cell lineage tree (upper) and the cell-segmented spatial view (lower). Shown are lineage trees with the four radially symmetric quadrants collapsed. “Z” - zygote. (**G**) RNA domain analysis identified transcriptomically distinct subcellular domains. The example embryo is visualized with its transcripts colored by cells (left), RNA domains (middle), and gene identities (right).

The mud snail, *Ilyanassa obsoleta*, a species of spiralia clade, is one of the most characterized systems of asymmetric mRNA inheritance. Work from the Lambert group estimated that 3–4% of embryonic RNAs show asymmetric localization within cells, and suggested that a subset of these transcripts are required for establishing cell lineage fates, including the eye (*14–16, 25*). However, these studies relied largely on nonquantitative, gene-by-gene *in situ* measurements in separate embryos. Hence a quantitative view of how multiple mRNA species are organized within each cell and partitioned across cell divisions is still lacking. Further, it remains unclear to what extent mRNA asymmetric segregation extends to other spiralians. We have recently developed the marine slipper snail, *Crepidula atrasolea*, as a laboratory model for spiralian development (*26*), providing a system in which to examine mRNA partitioning quantitatively across many transcripts and defined embryonic lineages.

Methodologically, we adapted a spatial transcriptomic technique, MERFISH (Multiplexed Error Robust Fluorescence In situ Hybridization), to examine mRNA localization and inheritance in *C. atrasolea*. MERFISH has been widely used to profile hundreds to thousands of genes at single-molecule resolution in mammalian tissue slices, as well as dissected whole zebrafish embryos (*27–29*). Here we have enhanced the 3D capabilities of MERFISH to accommodate whole-mount embryos for the first time. Using whole-mount MERFISH, we profiled over 200 genes at single-molecule resolution through the entire volume of the 350um thick *C. atrasolea* embryos. To complement the snapshot views from MERFISH, we developed a live reporter to track the dynamic spatial organization of total endogenous mRNAs in live embryos.

Using this approach, we characterize the emergence of transcriptomically distinct cell types during *C. atrasolea* early development with high temporal resolution, from the 1-cell through 24-cell stages across 43 embryos. We show that cell-type-specific transcriptomic profiles arise primarily through asymmetric partition of pre-existing mRNAs during mitosis, rather than through differential genomic regulation. We identify the subcellular domains mediating asymmetric mRNA partitions, reconstruct their cell-cycle dependent dynamic trajectory, and propose a quantitative model for transcript sorting at these domains that depends on transcript abundance and their affinities to the domains. Our study demonstrates that post-transcriptional mRNA localization can play major roles in cell type patterning and establishes an experimental and analytical framework for investigating mRNA localization at lineage, cellular and subcellular levels.

## Results

### 3D whole-mount spatial transcriptomics allowed multi-scale transcriptomic analysis at lineage, cell, and subcellular levels

We implemented 3D MERFISH to image the localization of single mRNA molecules across 216 genes in whole-mount *C. atrosolea* embryos. The 216 genes selected include classic developmental transcription factors (e.g. *tbxt*), signaling pathway components (e.g. *dl*), cell cycle regulators (e.g. *pcnt*), cytoskeletal components (e.g. *tba1a*), genes with known subcellularly localized transcripts (e.g. *aspm*), as well as genes that are highly and dynamically expressed in early *C. atrasolea* development according to a bulk RNA-seq dataset that included 12 embryonic stages from 2-cells to 53-cells (table S1). Probes targeting the selected genes were designed against our published *de novo* transcriptome assembly (Materials and methods, Data S1) (*30*).

Building on our previous developments of 3D MERFISH applied to Zebrafish development (*28*), we modified the methodology to enable higher imaging depth (up to 350 um) covering entire embryos. First, as before, we used acrydite-modified MERFISH probes targeting mRNAs. This allowed the covalent integration of the probes into the acrylamide gels embedding the embryos, preventing loss of probes during prolonged sample clearing and hybridization procedures. Second, instead of confocal microscopy, which limits the brightness of single molecules and prohibits imaging for depths >150um, epifluorescence imaging was employed using a high numeric aperture and large working distance objective. Crucially, we optimized our combinatorial MERFISH imaging by reducing the number of genes imaged in each cycle of hybridization to ensure that optical crowding was controlled for and that an epifluorescence strategy was feasible (Materials and methods). Third, to analyze the resulting imaging data with extended imaging depth, deconvolution was done with depth-dependent point spread functions; further, to combat the spatial distortions in the thick gel over multiple rounds of hybridizations, an iterative distortion correction and decoding pipeline was implemented to facilitate faithful decoding of single molecule identities (Materials and methods).

In total, 43 embryos at early cleavage stages from 1-cell to 24-cells were profiled by whole-mount 3D MERFISH (**Fig. 1B-C**). Decoded MERFISH recapitulated the expression patterns from single-gene smFISH and agreed with bulk RNA-seq data at corresponding stages (fig. S1A-C).

By sampling many embryos, we recapitulated the stereotyped spiral cleavage program within our MERFISH data (**Fig. 1B, D-E**) (*22, 31*). Briefly, the zygote undergoes two cleavages to form four large cells, termed “macromeres”, establishing the four “quadrants” of the embryo. Each macromere then divides twice asymmetrically, each time producing a smaller “micromere” daughter. This generates a 12-cell embryo organized in three tiers, or “quartets”, along the animal–vegetal axis, with 4 macromeres on the vegetal pole, and two quartets of micromeres on the animal pole. In the next three rounds of cleavages, cells in each quartet divide once, resulting in a total of 24 cells. From the 4- to 24-cell stages, embryos largely exhibit tetraradial symmetry across the four quadrants both morphologically and lineage-wise, with subtle cell-cell contact differences and cell cycle asynchronies that inform specific-quadrant identities.

We adopted a standard nomenclature to annotate cell lineage identities: uppercase and lowercase letters denote macromeres and micromeres, respectively; letters A–D or a-d specify quadrant identity; and Q or q refers to cells within a quartet without specifying quadrant identities (*16, 28*). A numeric prefix denotes the round of macromere cleavage that produced the cell, and superscripts 1 or 2 indicate animal or vegetal daughters (**Fig. 1D-E**). For instance, D gives rise to 1D and 1d, and 1d further divides into 1d1 and 1d2. Cell segmentation was performed using a modified neuronal network approach based on Cellpose (*32*), using both nuclear signal (DAPI), and MERFISH transcriptome signal (fig. S1D, Materials and methods). Segmentations were manually curated and cell lineages were annotated according to their embryonic positions and morphologies. This allowed gene expression analysis at single-cell resolution and temporally across cell lineages (**Fig. 1F**). Leveraging the single-molecule resolution of the MERFISH data, we have developed an RNA domain analysis pipeline to define the subcellular organization of RNA within the embryos. This approach identified transcriptomically distinct spatial domains in the embryo beyond single cell units, uncovering many subcellular domains with distinct RNA compositions (**Fig. 1G**).

In summary, we have combined whole-mount 3D MERFISH with a lineage-resolved embryonic system and novel RNA domain analysis to connect transcriptomic organization at single-cell, cell lineage, and subcellular levels. We have made the exploration of this data available within a custom browser at spiralia.merfisheyes.com.

### Transcriptomically distinct cell types corresponded to cell quartets in early C. atrasolea **embryos**

Although cell lineages and fates have been resolved through lineage tracing studies in many spiralians since the late 19th century (*19, 33*), molecular similarities and differences between embryonic cells remain poorly understood. Upon applying standard unsupervised clustering (entailing total count normalization, PCA, and Leiden clustering) to the MERFISH-derived single cell transcriptomes, we observed that embryonic cells across early cleavage stages formed transcriptomically distinct clusters, or cell types (**Fig. 2A**). These transcriptomically distinct cell types partially recapitulated embryonic lineage relationships: strong transcriptomic separation occurred across different quartets (**Fig. 2B-left, Fig. 2C**), as well as between polar bodies and blastomeres (**Fig. 2B-middle**). By contrast, quadrant identities did not separate in the transcriptomic space (**Fig. 2B-left, Fig. 2D**), suggesting that transcriptomic profiles of cells mirror their lineage and morphological tetraradial symmetry in the embryos.

**Fig. 2.**
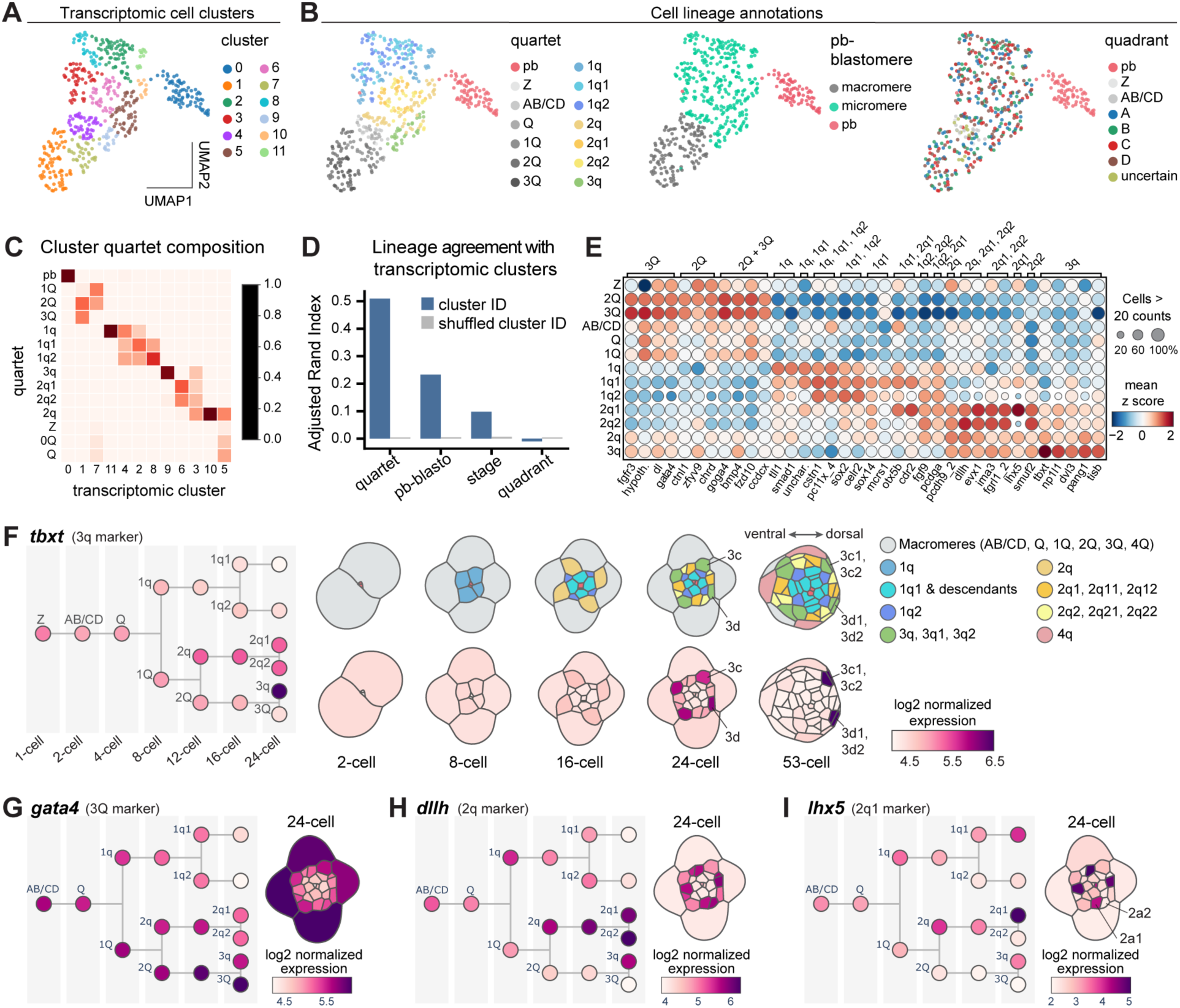
Cells from different quartets are transcriptomically different. (**A**) UMAP projection of single-cell transcriptomes colored by cluster identities. Transcriptomic clusters are identified by Leiden clustering. (**B**) The same UMAP projection with cells colored by lineage identities. “pb” - polar body; “Z” - zygote. (**C**) Heatmap showing fraction of cells in each transcriptomic cluster from each quartet. (**D**) Agreements between cell partitioning by transcriptomic clustering and various cell lineage groupings. Higher adjusted rand index indicates better agreement. Agreements between randomly shuffled cluster labels and lineage annotations are included for comparison. (**E**) Expression of quartet marker genes shown as a dot plot. “hypoth.” - hypothetical protein; “unchar.” – uncharacterized protein. (**F**) Expression of *tbxt* overlaid on quadrant-collapsed lineage tree (left) and cell-segmented embryos across stages (right). (**G-I**) Expression of *gata4*, *dllh*, and *lhx5* on lineage tree (left) and in an embryo at the 24-cell stage (right). Embryos are shown in animal views in **F-I**.

To identify marker genes that are enriched in specific lineages, we used a robust linear model to fit each gene’s single-cell expression as a function of cells’ quartet or quadrant identities. Genes that passed thresholds for model coefficient, significance, and expression fold-change were selected as markers (Materials and methods). This approach identified 36 marker genes across cell quartets, with some genes enriched in multiple quartets (**Fig. 2E**, fig. S2A-C, table. S2). No quadrant markers were found (fig. S2D), consistent with their lack of transcriptomic separation into distinct clusters. No enriched marker genes were found for early cell lineages before the 8-cell stage (Z, AB/CD, and Q), or for the 8-cell stage macromeres (1Q), whereas all other cell quartets were associated with at least one marker gene (**Fig. 2E**, fig. S2B). Among these markers, some are classic developmental regulators (e.g. *tbxt*, *gata4*), and many (e.g. *cstn1*, *hlx5*) have not been previously described in early spiralian embryogenesis (**Fig. 2E**, table. S2).

We highlight some of the quartet markers that are highly conserved developmental regulators. *tbxt* (*brachyury*) is a transcription factor broadly implicated in posterior mesoderm specification (*34, 35*). Our analysis identified *tbxt* as a marker for 3q (**Fig. 2F**), consistent with lineage tracing results that showed substantial contribution of 3q descendents to muscle tissues in the larva, including foot, body muscle, and esophagus (fig. S3) (*31, 36*). In many invertebrate and vertebrate embryos, GATA family transcription factors, including *gata4*, *gata5*, and *gata6*, function as endoderm specification factors (*37, 38*). Accordingly, we found *gata4* enriched in the macromeres (**Fig. 2G**), which are known endoderm progenitors (fig. S3). The 2q marker *dllh* is a homolog for *distal-less*, a highly conserved homeodomain transcription factor with well-described roles in neural development (**Fig. 2H**) (*39, 40*). In line with this, many larval neuronal cells are derived from the 2q lineage (fig. S3). Our analysis also identified markers that distinguish between lineages assigned with the same fate. For example, although the 2q daughter cells 2q1 and 2q2 were shown to contribute to neuroectoderm tissues within the same spatial domains in the larvae (*31, 36*), we found that 2q1 and 2q2 can be distinguished by the 2q1-enriched *lhx5* (**Fig. 2I**). *lhx5* is a transcription factor involved in neural patterning (*41, 42*), supporting the possibility that 2q1 and 2q2 give rise to molecularly distinct neuronal subtypes.

Cells in the four quadrants (A, B, C, and D) showed little transcriptomic difference between each other throughout 4- to 24-cells stages (**Fig. 2B, D**, fig. S2D) despite distinct lineage outcomes (fig. S3) (*31, 36*). This is highlighted in 3q, the third-quartet micromeres: although the 3a, 3b, 3c and 3d cells are transcriptomically similar, lineage-tracing showed that 3a and 3b contributed to esophagus and body muscle, whereas 3c and 3d contributed to foot muscle (fig. S3) (*36*). We therefore asked if quadrant-specific expression patterns emerge after the morphological tetraradial symmetry of the embryo breaks. Indeed, at the 53-cells stage, shortly after the D quadrant macromere 3D adopts a cleavage program distinct from the other quadrants to establish the dorsal side of the embryo, we detected transcriptomic differences among quadrants. For example, expression of the 3q marker *tbxt* was maintained in the 3c and 3d descendants (foot muscle progenitors), which are on the dorsal side of the embryo, but was missing in the 3a and 3b descendants (esophagus and body muscle progenitors) on the ventral side (**Fig. 2F**).

In summary, by combining MERFISH derived single-cell transcriptomes with lineage annotations, we showed that cells in early cleavage differ not only in their future fates, but also in their transcriptomic profiles. Transcriptomically different cell quartets can be observed as early as the 8-cells stage, after the first embryonic cleavage along the animal-vegetal axis. Each subsequent cell cleavage created additional transcriptomically unique cell quartets. In contrast, transcriptomic patterning of quadrants along the dorsal-ventral axis was not observed until the axis became morphologically apparent. Taken together, the molecular patterning of the animal-vegetal axis precedes dorsal-ventral axis and is coupled with early cleavage events.

### Lineage-specific transcripts undergo subcellular aggregation and asymmetric partition

Several lines of evidence suggested that zygotic transcription is low in early *C. atrasolea* embryos until after the 24-cell stage. First, our FISH assay revealed that nuclei at early cleavage stages exhibited low levels of total mRNA (polyA), rRNA (18S), and the spliceosome snRNA (U6) (fig. S4). In contrast, embryos at and past the 53-cell stage exhibited nuclear enrichment of these RNAs, indicative of active zygotic transcription at these later stages (fig. S4). Second, small cytoplasmic granules enriched with mRNAs appeared transiently between the 53-cells stage and the onset of gastrulation (fig. S4D), consistent with degradation granules associated with maternal mRNA turnover during the maternal-to-zygotic transition (MZT) in other species (*43, 44*). These observations suggest that cell quartets in early embryos acquired differences in their mRNA composition when zygotic transcription was not fully active.

To understand how transcriptomic diversity arose with low active transcription, we visualized gene expression patterns at single-molecule level across developmental stages (**Fig. 3A-D**, fig. S5). Transcripts of many lineage markers were distributed unevenly within cells, with markers of daughter cells enriched in specific regions of their mothers before division (**Fig. 3A-D**, fig. S5). We highlight the example of the first quartet micromeres (1q) at the 8-cell stage, which are daughters of the 4-cell stage macromeres (Q). Before the birth of 1q, transcripts of the 1q marker *cstn1* were enriched on the animal side of the dividing Q cell (**Fig. 3B**). Similarly, the 2q marker *dllh* and the 3q marker *tbxt* formed subcellular aggregation in 1Q (2q mother) and 2Q (3q mother), respectively (**Fig. 3C-D**).

**Fig. 3.**
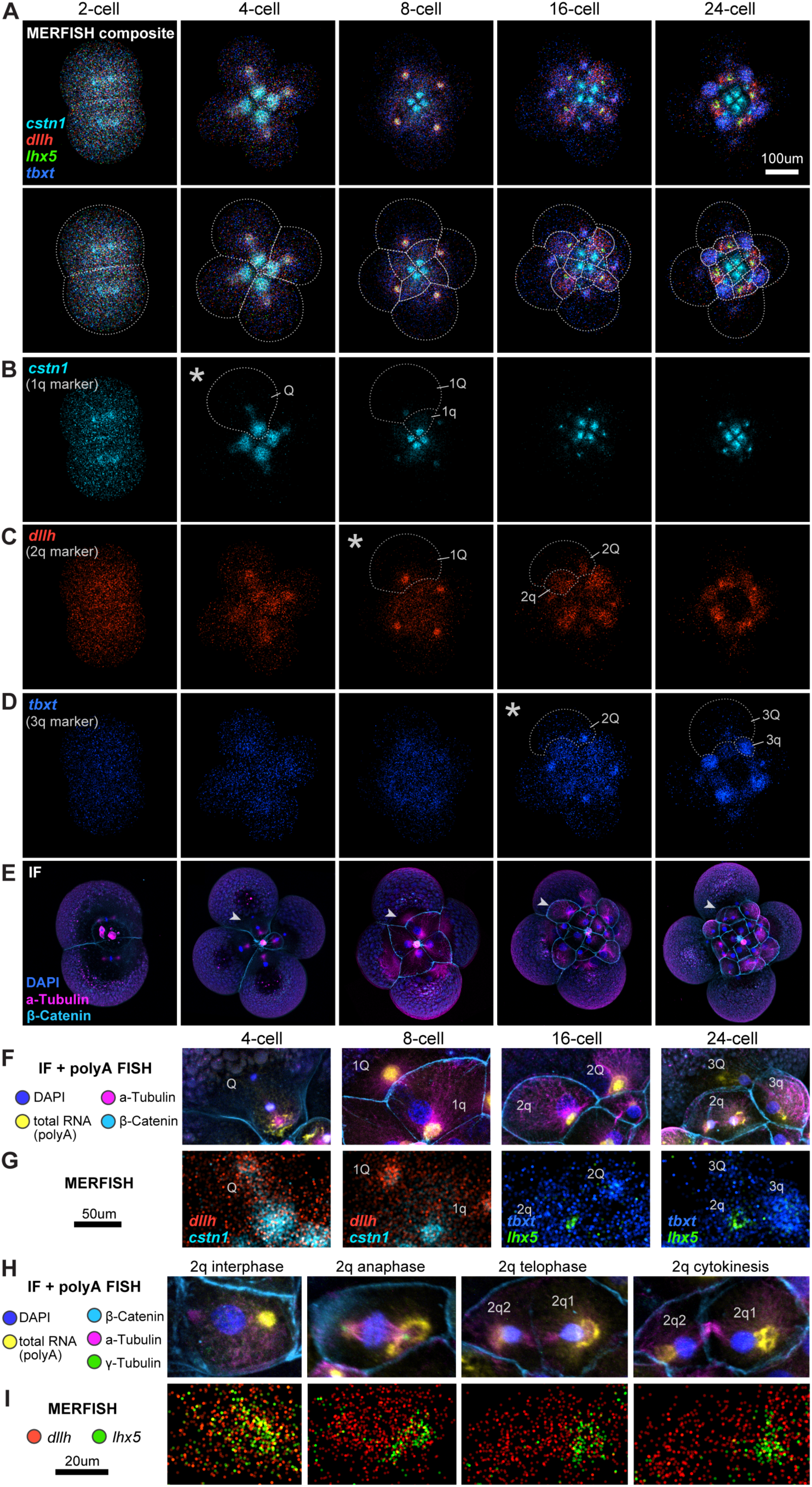
Transcript distribution at single-molecule resolution reveals their subcellular aggregation. (**A-D**) MERFISH of selected marker genes across developmental stages. Points correspond to single transcripts colored by gene identities. Animal projections of embryos are shown. Cells are outlined with dashed curves in **A** (lower panel). Asterisks in **B-D** mark the stages each gene first displays strong aggregation. Dashed curves in **B-D** outline example cells with transcript aggregation and their daughter cells in the next stage. (**E**) Immunofluorescence on the same embryos as in **A-D**. Arrowheads point to the regions enlarged in **F-G**. (**F**) Enlarged insets from **E** with the addition of PolyA FISH for total mRNA distribution. (**G**) Distributions of selected marker transcripts in the same insets as in **F**. (**H-I**) Second quartet micromeres (2q) across cell cycle phases. **H** and **I** show polyA FISH-coupled immunofluorescence and MERFISH, respectively, in the same cells. *lhx5* transcripts are asymmetrically partitioned into the 2q1 daughter cell.

To visualize how subcellularly aggregated transcripts become enriched in selected daughter cells, we combined immunofluorescence with 3D-MERFISH in the same embryos. Antibodies against beta-catenin and alpha-tubulin labeled cell membrane and microtubules, respectively, which helped define the subcellular structures transcripts were associated with and the cell cycle phase the aggregates appeared in (**Fig. 3E-F**). This revealed that, during interphase, transcripts of marker genes were surrounded by microtubule arrays (**Fig. 3F-G**, 8-cell, 16-cell). In contrast, during anaphase, marker transcripts were concentrated at one of the mitotic spindle poles corresponding to the daughter cell they mark (**Fig. 3F-G**, 4-cell, 24-cell).

The highly stereotypic development of the *C. atrasolea* embryo enabled the assembly of a pseudo-timecourse from different cells of the same quartet identity at progressing cell cycle phases (**Fig. 3H-I**). Such pseudo-timecourse revealed how transcripts became asymmetrically partitioned between the daughter cells through mitosis. For instance, during 2q division, *lhx5* transcripts showed increasing restriction around the mitotic spindle pole at the 2q1 daughter side. This resulted in the majority of *lhx5* transcripts being inherited by the 2q1 daughter and, correspondingly, their depletion from the 2q2 daughter upon cytokinesis (**Fig. 3H-I**). Similar asymmetric aggregation and partition was widely observed for other marker genes (**Fig. 3A-D**, fig. S5).

Taken together, integrating immunofluorescence and 3D MERFISH, our results suggest that, in contrast to classic examples of cell type specification, transcriptomically diverse cell types in *C. atrasolea* are created largely independent of transcriptional regulation. Instead, subcellular aggregation and asymmetric partition of specific transcripts during mitosis drives the simultaneous diversification of cell lineages and their molecular contents.

### mRNAs exhibit dynamic centrosome and cell cortex associations at early cleavage stages

Our observations of asymmetrically partitioned mRNAs during mitosis raise the possibility that transcripts are organized around the mitotic apparatus. To test this possibility, we sought to visualize the dynamic interplay between mRNAs and mitotic apparatus with high temporal resolution. Total mRNA, marked by polyA FISH, was concentrated at the sites of marker gene aggregation (**Fig. 3F, H**). This suggests that many transcripts follow similar trajectories to asymmetrically segregate into specific cell lineages. We therefore performed live imaging using a total mRNA reporter to visualize the spatiotemporal trajectory of mRNA aggregation and partitioning.

We developed a total mRNA reporter by constructing a fluorescently tagged polyA-binding protein, PABP(N)-mNeonGreen. This reporter contains the N-terminus of the human polyA-binding protein, PABPC1, which harbors four RNA recognition motifs that bind mRNAs at polyA tails (**Fig. 4A**). Embryos were injected with mRNAs encoding the reporter at the 1-cell stage and imaged from the 4-cell stage (**Fig. 4B**). Fluorescent reporters for nuclei (H2B-CFP), cell cortex (LifeAct-CFP), and microtubule (EMTB-mRuby) were added to provide cellular context (Materials and methods). The resulting movie captured the cell-cycle-dependent localization of the mRNA aggregation domains, consistent with observations from the MERFISH snapshots (**Fig. 4C**, movie S1).

**Fig. 4.**
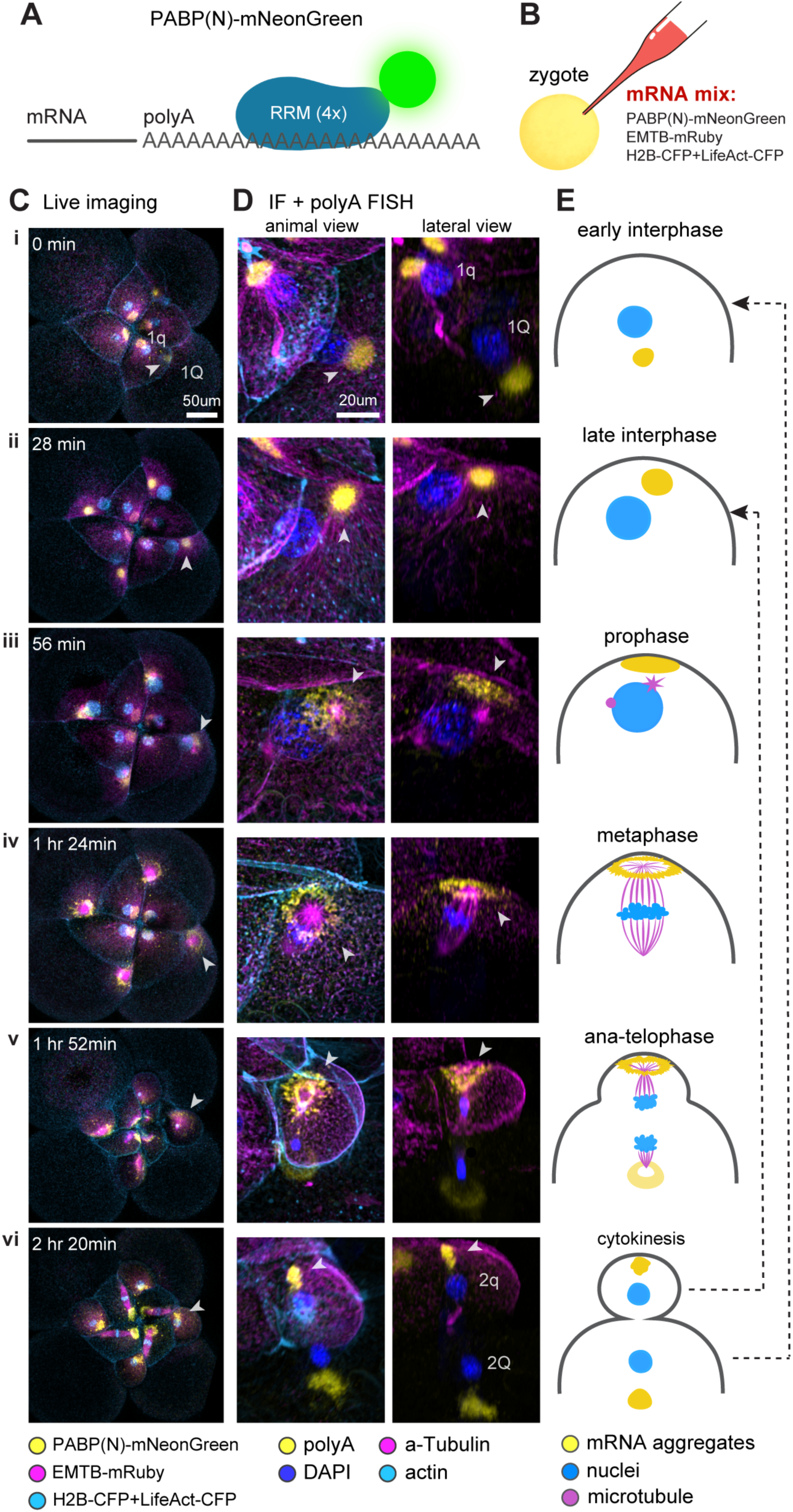
Live imaging and polyA FISH-coupled immunostaining reconstruct the spatiotemporal trajectory of mRNA aggregation. (**A**) Schematic of the live total mRNA reporter, PABP(N)-mNeonGreen. “RRM” - RNA recognition motif. (**B**) Embryos were injected with mRNAs encoding the reporters at the zygote (1-cell) stage for live imaging. (**C**) Frames from live imaging of an embryo undergoing 1Q mitosis (8- to 12-cell stage transition). The full movie spanning the 4-cell to 42-cell stages with 7-min intervals is available as movie S1. (**D**) PolyA FISH-coupled immunostaining of embryos stage-matched to the ones in **C**. Shown are insets focusing on one dividing macromere per embryo. Arrowheads point to the “same” mRNA aggregates in **C** and **D** based on their subcellular location and shape. mRNA aggregates marked by arrowheads fall on the same continuous spatial trajectory across time according to the live movie. (**E**) Illustrations of the observed mRNA aggregation trajectory throughout mitosis. Dashed arrows mark the starting points of daughter cell trajectories: the vegetal side daughter cell repeats the same trajectory, while the animal side daughter cell repeats the trajectory from the “late interphase” configuration.

Live imaging provided high temporal resolution, but signal diminished as imaging depth extended into the yolky macromeres. Hence we complemented the live imaging with polyA-FISH-coupled immunostaining across 40 additional fixed embryos. A refractive-index-matching medium (BBBA) allowed imaging of structures deep inside the macromeres. Leveraging the highly stereotypical early developmental program, we matched the immunostained embryos to frames in the live imaging movie (**Fig. 4C-D**), providing two complementary approaches for tracking mRNA aggregation across time.

We found that mRNA aggregation iterated a similar trajectory during each cell cycle (**Fig. 4E**). In macromeres, such as 1Q, a primary mRNA aggregate appeared as a globule at the early interphase on the vegetal side of the nucleus (**Fig. 4D-E i**). This primary aggregate grew in mRNA density and moved to the animal side of the cell just below the cell cortex by late interphase (**Fig. 4D-E i-ii**). As cell cycle progressed into prophase, the globular aggregate appeared flattened against the cell cortex and slightly dispersed (**Fig. 4C-E iii**). Meanwhile, two centrosomes formed, with one positioned under the mRNA aggregate domain (**Fig. 4C-E iii**). At metaphase, the mitotic spindle was established and astral rays could be seen around the animal side centrosome (**Fig. 4C-E iv**, fig. S6A). Concomitantly, the primary mRNA aggregate formed a ring-like structure centered at the centrosome (**Fig. 4C-E iv**, fig. S6A). During anaphase and telophase, a secondary, smaller mRNA aggregate formed as a ring around the vegetal side spindle pole (**Fig. 4C-E v**). During cytokinesis, the mRNA ring at each spindle pole retracted (**Fig. 4C-E vi**). After the cell division completed, the secondary mRNA aggregate inherited by the vegetal daughter became the new globular primary aggregate in the cell, and the same trajectory described above repeated (**Fig. 4E**, fig. S6B). In the animal daughter cell, mRNAs recollected into a domain below the cell cortex, reminiscent of the primary aggregate at the late interphase in the mother cell (**Fig. 4E**, fig. S6B). This mRNA configuration persisted through the interphase of the animal daughter cell, after which the described trajectory resumed (**Fig. 4E**, **Fig. 3H**).

The observed mRNA trajectories revealed an interplay between mRNA and microtubule structures. During late interphase, mRNA aggregate colocalizes with a microtubule organizing center (MTOC), which is marked by gamma-tubulin and connected with microtubule arrays (**Fig. 4D ii**, fig. S6C). During metaphase to telophase, mRNAs appear to associate with the astral microtubules centered around the gamma-tubulin-labeled centrosome (**Fig. 4D iv-v**, fig. S6A). This suggests that mRNA localization may be dependent on microtubule structures. Corroborating this idea, pharmacological disruption of microtubule dynamics by nocodazole interrupted both microtubule and mRNA spatial organization and caused cell cycle arrest (fig. S6D), similar to previous observations in the mud snail (*14*). In contrast, inhibition of actin filaments by cytochalasin-D left mRNA trajectory largely unaffected despite the abolishment of cytokinesis (fig. S6E).

In summary, mRNAs in early *C. atrasolea* embryos follow a cell-cycle- and microtubule-dependent localization program. This localization program is gradually abolished after the 24-cells stage, and by the 53-cells stage, cells stopped forming MTOC or cell-cortex-associated mRNA aggregates (movie S2, fig. S4D). The abolishment of the mRNA aggregation program coincides with the increase of zygotic transcription, as measured by the nuclear RNA content (fig. S4). This suggests that mRNA aggregation and asymmetric partition may be a mechanism required for establishing molecularly different cell types when zygotic transcription is not fully available.

### An affinity-competition model predicts marker gene sorting at subcellular aggregates

While total mRNA tracking highlights the dynamics of mRNA aggregate formation and inheritance, MERFISH makes it possible to quantify gene composition within the aggregates. Marker gene distribution (**Fig. 3A-D**, fig. S5) suggests that aggregate composition varies across cell lineages. For instance, the 1Q aggregates exhibit a high density of the 2q marker *dllh*, while the 2Q aggregates display a high level of the 3q marker *tbxt* (**Fig. 3C-D**).

To systematically identify the mRNA aggregates and characterize their transcript compositions in the MERFISH data, we developed an RNA domain analysis. Our approach does not depend on pre-defined cellular or subcellular segmentations. Instead, we first group spatially proximal transcripts into communities (350 nearest neighbor transcripts), then cluster these communities based on their transcriptomic profiles, resulting in RNA domains that are internally coherent in gene composition, but distinct from each other (**Fig. 5A**, Materials and methods).

**Fig. 5.**
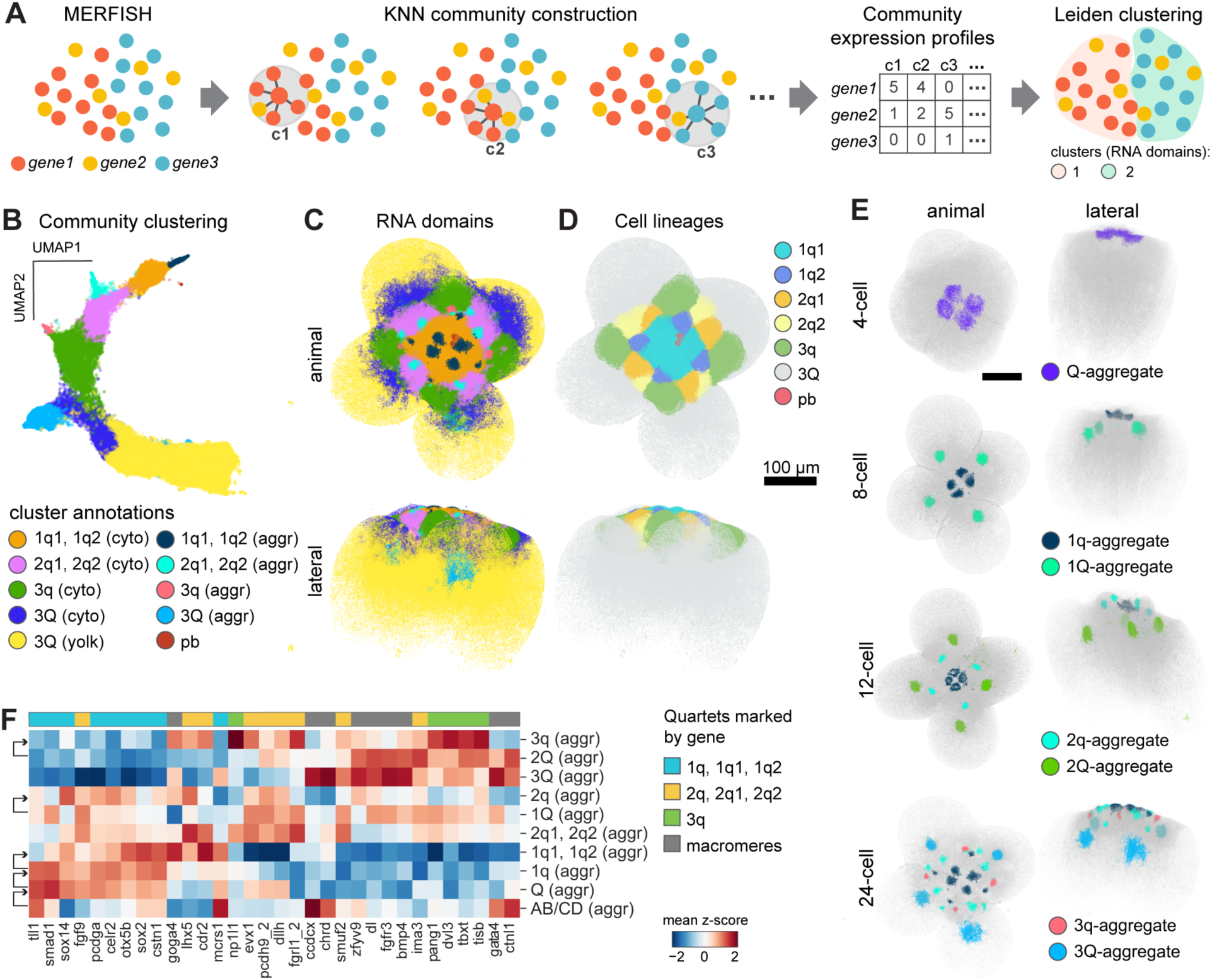
Characterization of gene composition at subcellular aggregates. (**A**) Schematic of the RNA domain analysis. “c1”, “c2”, “c3” - community 1, 2, 3. (**B**) UMAP projection of the communities from the example embryo colored by annotated Leiden clusters (RNA domains). “Cyto” - cytoplasm; “aggr”: aggregate. (**C**) Spatial view of all transcripts in the example embryo colored by annotated RNA-domains. The same coloring scheme in **B** is used. (**D**) The same embryo with transcripts colored by cell quartet identities. (**E**) Embryos across developmental stages with transcripts in aggregate domains highlighted. All other transcripts are shown in grey. (**F**) Heatmap showing the gene expression z-score of the previously identified quartet markers across aggregate domains. Rows and columns are ordered by hierarchical clustering. Arrows on the left indicate mother to daughter relationship between cells represented by adjacent rows. The colorbar on top indicates which quartet each gene marks. If a gene is a marker of multiple quartets, the color shows the quartet with the largest expression fold change.

Applying this approach to a representative 24-cell stage embryo, we identified 10 RNA domains annotated according to their cellular and subcellular identities (**Fig. 5B-C**). As expected, transcriptomically diverse cell quartets separated as different domains (**Fig. 5B-D**). Further, subcellular aggregates were identified as distinct domains from their corresponding cell cytoplasm, reflecting compositional differences between the aggregates and the remaining cytoplasm (**Fig. 5B**). We performed the RNA domain analysis on all embryos and extracted the domains corresponding to the subcellular mRNA aggregates (**Fig. 5E**). Quartet marker gene expression confirmed that aggregate composition varies across cells (**Fig. 5F**).

How do different cells select different groups of transcripts to join the aggregate? In one model, the mRNA aggregation site exhibits cell-type-specific transcript affinities. For example, differences in the RNA-binding proteins present at aggregates across cells could result in different transcript preferences. Alternatively, we propose a simpler model, where aggregates exhibit transcript-specific affinities that are fixed across cells, while transcripts compete for limited binding capacity. In this scenario, high-affinity transcripts would aggregate and become depleted first, enabling lower-affinity transcripts to accumulate in descendent cells for subsequent cell divisions (**Fig. 6A**). We term this model the fixed-affinity-competition model, or simply affinity-competition model.

**Fig. 6.**
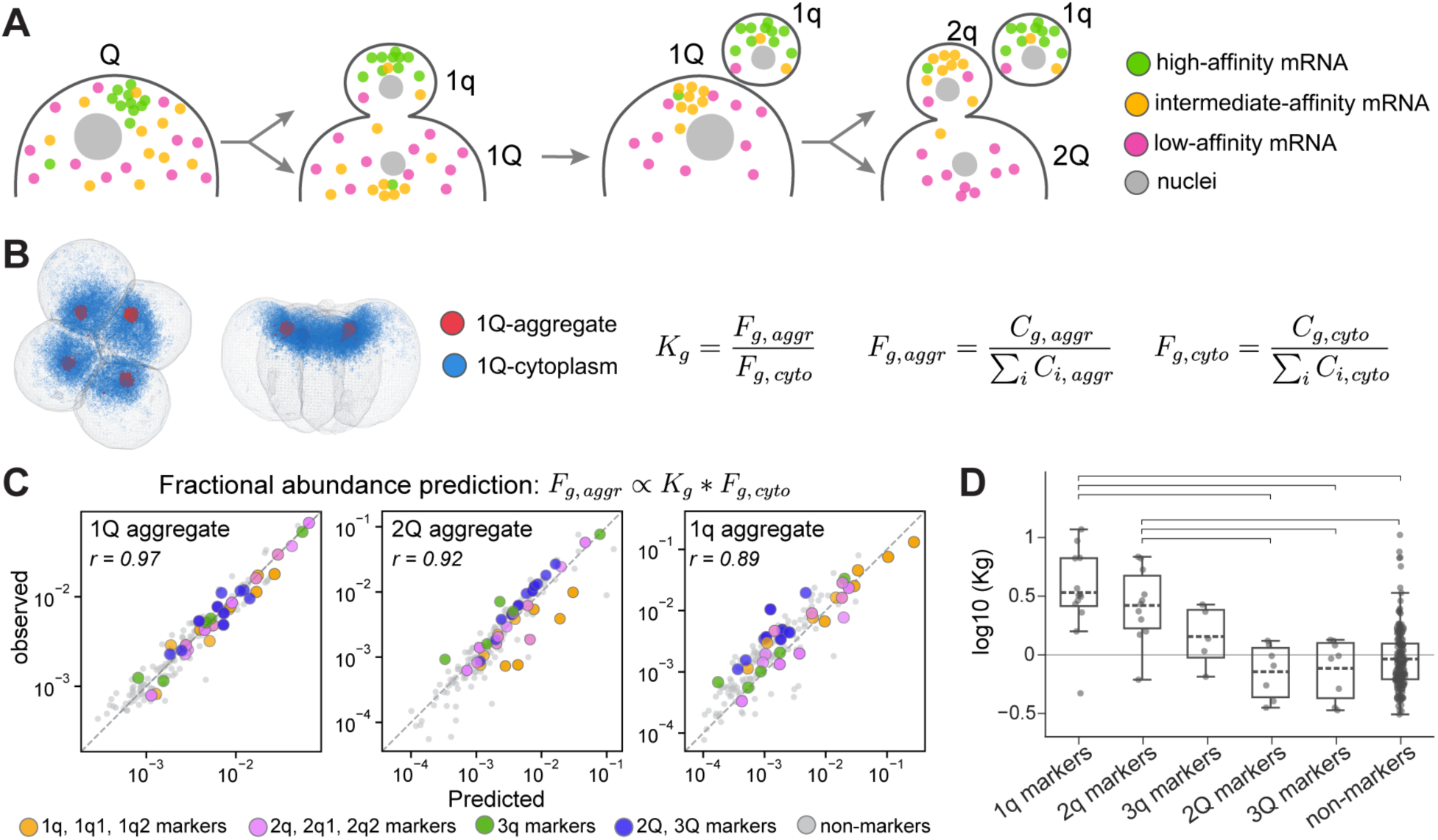
An affinity-competition model predicts gene composition at subcellular aggregates. (**A**) Schematic of an affinity-competition model that generates transcriptomically diverse cells. (**B**) Formulation of the affinity-competition model. Left: transcripts in the 1Q aggregate and the surrounding cytoplasm are shown in red and blue, respectively, in an 8-cells stage embryo. Right: Formulae for calculating Kg, the affinity coefficient of gene g, and the fractional abundances used in Kg calculation. “F” - fractional abundance. “C” - counts. Subscripts: “g, aggr” - transcripts of g in the aggregates; “g, cyto” - transcripts of g in the cytoplasm. (**C**) Observed and predicted gene composition at subcellular aggregates. Kg inferred from 1Q and Fg, cyto observed in the corresponding cells are used for the predictions. (**D**) Comparison of Kg values for different marker gene groups. 1q markers include 1q, 1q1, and 1q2 markers. 2q markers include 2q, 2q1 and 2q2 markers. Genes that mark multiple quartets are replotted for each group. Boxes show the interquartile range (25th-75th percentile) with dashed lines marking the mean. Whiskers extend to the most extreme data points within 1.5x the interquartile range. Brackets indicate groups with adjusted p-values below 0.01 from a Mann–Whitney U test.

To test whether aggregate gene composition in different cells can be explained by the affinity-competition model, we leveraged the quantitative nature of the MERFISH data and performed mathematical modeling. We formulated the affinity-competition model based on the extended Langmuir adsorption model (*45, 46*). Under this framework, transcripts associate with the aggregation domain according to their cytoplasmic concentration and a transcript-specific affinity coefficient. Assuming mRNA aggregation domains are saturated at equilibrium, the relative affinity coefficient (Kg) for a given gene can be calculated as the ratio of its fractional abundance in the aggregate (Fg,aggr) to its fractional abundance in the rest of the cytoplasm (Fg,cyto). An affinity coefficient greater than 1 therefore indicates an enrichment of the transcript in the aggregate relative to the remaining cytoplasm. The fractional abundance of a transcript in the aggregate can in turn be estimated given its affinity coefficient and its fractional abundance in the cytoplasm (**Fig. 6B**).

Using this model, we first inferred transcript-specific affinity coefficients from the transcript abundances in the 1Q aggregation and cytoplasm domains (**Fig. 6B**). We next asked whether the same set of affinity coefficients can be used to predict aggregate compositions across different cells. With the affinity coefficients computed from 1Q and the observed cytoplasmic transcriptomic profiles, we predicted the aggregate transcript compositions in 1q and 2Q. Predicted and observed aggregate compositions were highly correlated (**Fig. 6C**). Therefore, a common set of transcript-specific affinities can explain aggregate compositions across cells without invoking cell-type-specific aggregation mechanisms.

The affinity-competition model further predicts that transcript-specific affinities determine the differential mRNA profiles across cells based on their birth order. High-affinity transcripts are recruited first and segregated into the earliest-born micromeres, whereas their depletion from the daughter macromere frees aggregate capacity for lower-affinity transcripts during subsequent divisions (**Fig. 6A**). Accordingly, 1q markers should exhibit the highest affinities, followed by 2q and 3q markers, while macromere markers and non-marker genes should have the lowest affinities. Mean affinity coefficients estimated across all macromeres supported this hierarchy: 1q and 2q markers displayed the highest affinities, whereas macromere markers and non-marker genes showed significantly lower affinities (**Fig. 6D**).

In summary, we modeled gene sorting in the mRNA aggregates with an affinity-competition model. Further, cell-type-specific aggregate composition does not require cell-type-specific aggregation mechanisms. Instead, early aggregation and depletion of high-affinity transcripts progressively reshapes the cytoplasmic transcript pool, enabling the progressive segregation of distinct transcript populations across cell quartets.

## Discussion

In this study, we implemented a state-of-the-art 3D MERFISH platform to profile gene expression across hundreds of genes in whole-mount embryos measuring 350 um. This provides a cost-effective solution for direct volumetric spatial transcriptomics imaging in which expensive confocal or lightsheet instrumentation can be replaced by inexpensive high-numerical-aperture and long-working-distance epifluorescence instrumentation. Critically, this methodology allowed us to generate the first lineage-resolved single-cell level transcriptomic analysis of spiralian embryos, for which single-cell sequencing assays are typically not feasible due to their difficulties to dissociate. We also engineered a live imaging system for tracking the subcellular localization of total mRNA during embryogenesis. Using a combination of whole-mount MERFISH, immunofluorescence, live imaging, and subcellular RNA domain analysis, we show that, in early *C. atrasolea* embryo, specific transcripts concentrate at mitotic apparatus-associated aggregates, and are asymmetrically inherited by daughter cells during mitosis. Our methodology is applicable to other developmental systems where RNA subcellular distribution is understudied. Our findings expand our understanding of the developmental logic by which cellular diversity can be generated: cleavage geometry, cell-cycle dynamics, and RNA aggregation affinity together can convert a shared maternal transcriptome into molecularly distinct cell lineages.

Recent efforts in spatial transcriptomics have begun to connect molecular state, spatial context, and developmental progression by profiling thick tissues across developmental time. For example, weMERFISH profiled yolk-removed and flattened whole zebrafish embryos from gastrulation to organogenesis (*28*), while 3DEEP profiled intact mouse skin containing hair follicles spanning organogenesis stages (*47*). A challenge in such 4D spatial transcriptomic approaches is the reconstruction of the developmental progression: because transcriptomic profiling is destructive, lineage relationship between cells sampled at different developmental stages can only be inferred by molecular similarity and spatial distribution at the population level (*28, 47*). In contrast, leveraging the stereotyped geometry of spiral cleavage, our study enabled comprehensive lineage-resolved 4D spatial-temporal measurements. In whole-mount *C. atrasolea* embryos, cell lineage identities can be assigned unambiguously from embryonic position and morphology, rather than inferred from transcriptomic similarity. Our data demonstrates that transcriptomic similarities do not always agree with lineage relationships: in *C. atrasolea* embryos, a cell is often less transcriptomically similar to its sister cell (e.g. 1a and 1A) than to its corresponding cousin from a different quadrant (e.g. 1a and 1b). With dense sampling during early cleavage stages across 43 embryos, our approach allowed tracing transcript distributions through known lineages at single-transcript, single-cell, and single-division resolution.

The generation of diverse cell types during development is most often attributed to transcriptional regulation, where signaling pathways and transcription-factor networks produce cell-type-specific transcriptomic profiles *de novo* by transcribing different genes in different cells (*8, 9*). Here we show that, rather than relying on transcriptional regulation, cells can asymmetrically partition existing mRNAs to simultaneously produce new cells and new molecular identities. Using this strategy, *C. atrasolea* achieves 6 transcriptomically distinct cell quartets in a 24-cells stage embryo, highlighting asymmetric mRNA partition as an effective way of patterning a small cell population. Differential mRNA partitioning has been described in other developmental contexts (*17, 18*). In ascidian mesendoderm, *Not* mRNAs are partitioned into the mesoderm-fated daughter cells (*48*); during *Drosophila* neuroblast cell division, *prospero* and *inscuteable* mRNAs are asymmetrically partitioned into the ganglion mother cell and the self-renewing neuroblast, respectively (*49, 50*); and in the mud snail, over a dozen RNA species are differentially segregated into particular cell lineages by the 24-cells stage (*16*). Our study extends these examples using quantitative, highly-multiplexed methods and shows that asymmetric mRNA partitions can operate as a broad lineage-patterning mechanism. Virtually all micromere quartet marker genes we identified, ranging from transcription factors to adhesion molecules, were asymmetrically partitioned through subcellular aggregation. We further leveraged the single-molecule resolution of MERFISH data and developed a RNA domain analysis to characterize transcriptomically distinct spatial compartments without *a priori* segmentations.

While most spatial transcriptomic studies have focused on cell type and tissue domain analysis (*51, 52*), we applied a single-molecule level KNN approach to resolve subcellular RNA domains. This identified the subcellular RNA aggregates that mediate marker gene segregation and allowed the quantitative modeling of their compositions. Whereas previous work in the mud snail proposed intrinsic differences between centrosomes as the reason for differential accumulation of particular mRNAs (*14*), our mathematical modeling demonstrated that differential mRNA accumulation can occur even across centrosomes exhibiting the same mRNA binding properties. Together with our live imaging data, which revealed a temporal asynchrony between mRNA accumulation at the two centrosomal domains in the same cells, our model proposes that mRNAs with higher affinities are aggregated and depleted first, shifting the landscape of remaining mRNAs available for the next aggregate formation and partition.

Our study provides a framework for further exploring the mechanisms of RNA subcellular organization by identifying which specific RNA sequences are required for the differential localization of RNAs at subcellular domains. RNA localization is classically known to be regulated by *cis*-acting elements, often in the 3′ untranslated region (3’ UTR) of RNA (*53–55*). For example, the *β-actin* transcript in fibroblast contains a 54-nucleotide 3′ UTR ‘zipcode’ is bound by the ZBP1 protein; the complex is then carried along microtubules to cell periphery for local translation (*56–58*). In the mud snail, a 3′ UTR sequence from a localized mRNA (*IoLR5*) was necessary and sufficient for its centrosomal localization and asymmetric inheritance (*25*).

Our study suggests that, using 3D spatial transcriptomics, the subcellular distribution of endogenous transcripts or reporters can be quantitatively measured in high throughput *in situ*, allowing characterization of location-specific *cis*-elements. For instance, *C. atrasolea* transcripts could contain ‘zipcodes’ that mediate their affinity to the subcellular mRNA aggregation sites. As we scale 3D MERFISH to thousands of genes, hence obtaining more comprehensive measurement of transcript affinities to various subcellular locations, it will become possible to computationally predict location-specific zipcodes, which can then be tested by massively parallel reporter assays using MERFISH as the readout. Spiralian embryos present a unique model for such studies due to their abundant localized transcripts. Natural variation across spiralian species would further strengthen this framework. Our preliminary observations in the nudibranch *Berghia stephanieae* revealed localized mRNAs at the vegetal cortex rather than at MTOCs as in mud snail, slipper snail, and surf clam (*14, 59, 60*). Combining diverse spiralian models with expanded spatial transcriptomics will therefore facilitate the investigation of not only the *cis-* mechanisms underlying RNA localization, but also how localization programs are evolutionarily diversified.

Our findings add to growing evidence that molecular asymmetries during cell division are more widespread than previously appreciated (*17, 61–63*). We show that, in *C. atrasolea*, the overt geometric asymmetry of early cleavages is accompanied by a molecular asymmetry, in which MTOC-associated subcellular domains mediate the unequal partition of mRNAs between daughter cells. Since MTOCs organize cell polarity and spindle position, they are well-positioned to serve as platforms for asymmetric partitioning of molecular cargos (*13, 14, 17, 18, 64*). Notably, MTOC asymmetries have also been observed in cells previously considered to divide symmetrically (*62, 65, 66*). However, RNA distributions have not been systematically characterized in many of these contexts, raising the possibility that MTOC-mediated RNA partitioning is a broader and underappreciated mechanism for developmental patterning. With the framework demonstrated in this study, RNA localization can be characterized across a wide range of developmental contexts, opening new avenues for investigating the interplay between RNA localization, cytoskeletal organization, and developmental outcomes.

Spiral cleavage is considered the ancestral mode of early development for the spiralia clade (*24*), whose lineages radiated rapidly around 550 million years ago to span half of extant bilaterian phyla (*21, 67–69*). Our results and previous observations in other spiralian species raise the possibility that mRNA sorting is conserved across spiralians. If so, changes in which transcripts are sorted, and into which cells, are candidates for how spiralian embryos diversified, including the loss of spiral cleavage in lineages such as cephalopods (*21, 24, 70*). Comparing mRNA sorting across spiralians is thus a direct way to ask how modifications of one ancestral program accompanied the diversification of a large fraction of animal life.

## Acknowledgments

We thank members of the Bintu lab for helpful discussions, the Hamdoun lab for sharing their microscopes, and the UCSD School of Medicine Microscopy Core. We thank V. Barone, J. A. Gagnon, A. F. Schier for their feedback on the manuscript.

## Authors contributions

Conceptualization: Y.W., B.B., D.C.L.; Data Curation: Y.W., S.N., B.B.; Formal analysis: Y.W., B.B.; Funding acquisition: D.C.L. B.B.; Investigation: Y.W., T.P., E.I.; Methodology: Y.W., B.B.; Project Administration: Y.W., B.B., D.C.L.; Resources: Y.W., B.B., D.C.L.; Supervision: D.C.L., B.B., Y.W.; Visualization: Y.W., I.J.; Writing – original draft: Y.W., B.B., D.C.L.; Writing – review & editing:: Y.W., B.B., D.C.L..

## Data, code, and materials availability

MERFISH data, Immunostaining and live imaging data, as well as code used for data processing and reproducing results in this study have been deposited in Dryad (DOI: 10.5061/dryad.dz08kpsdn) and will be made publicly available upon publication of the manuscript. MERFISH data is additionally available for visualization through spiralia.merfisheyes.com. Plasmids generated in this study have been deposited at Addgene. Animal (*Crepidula atrasolea*) samples will be available upon request.

