## Supplementary Materials for "3D MERFISH reveals cell-type diversity generated by mRNA partitioning in spiralian embryogenesis"

**The PDF file includes:**

Materials and Methods
Figs. S1 to S6
Tables S1 and S2
Captions for Data S1
Captions for Movies S1 and S2
References (71–82)

**Other Supplementary Materials for this manuscript include the following:**

Tables S1 and S2 (.csv)
Data S1 (.fasta, with header map .csv)
Movies S1 and S2 (.mp4)

### Materials and Methods

*Crepidula atrasolea*

A breeding colony of adult wild-type *C.atrasolea* animals have been maintained since 2018 in the Lyons lab. Animals were kept at 27°C in 10cm petri-dishes with approximately 4 females and 3 males per dish on average. Animals were kept submerged in artificial sea water (ASW) made from Reef Crystals (RC1-200, Instant Ocean) and fed daily by diluting Phytofeast (250576, Reef Nutrition) in ASW at approximately 1:500 dilution. Dishes were cleaned daily by replacement of liquid content. Adult females were examined regularly for the presence of egg masses. Egg masses were collected by dislodging the female from the dish, flipping them such that the ventral side was up and egg masses were exposed, and transferring the egg cases to petri-dishes pre-filled with ASW containing antibiotics (penicillin 0.06g/L, streptomycin 0.2g/L) using forceps. Embryos were raised in their egg cases at room temperature (22°C) or at 27°C. Dishes were checked at least every 2 days to remove egg cases with high mortality rate and to exchange for fresh ASW with antibiotics until embryos reach desired stages.

Coating of dishes and pipette tips for embryo handling

Since both live and fixed embryos tend to stick to plastic petri-dishes, pipette tips, and glasswares, all labwares in direct contact with embryo samples need to be coated with gelatin or Bovine Serum Albumin (BSA), with the exception of the eppendorf lo-Bind tubes (022431048, 022431102, eppendorf), which exhibited low stickiness to fixed embryos. A 5x gelatin coating stock was made by dissolving 0.25g gelatin (Knox) in 50mL distilled water under heat followed by adding 600uL 16% formaldehyde solution when the gelatin mixture cools. The 5x gelatin stock was diluted in distilled water to 1x working coating solution. BSA coating solution was made fresh in distilled water at 0.5% BSA concentration. To coat a petri-dish, coating solution was poured into the dish, swirled to ensure even coating, then poured out. The coated dish was dried upside down on a paper towel. To coat a pipette tip (plastic or glass pasteur), the tip was attached to a pipette, the coating solution was pipetted up and down 3 times then fully dispensed. The tip was then detached from the pipette and placed tip up on a rack to dry.

Embryo decapsulation

Each egg case, or capsule, encases approximately 10 naked embryos. To release embryos from their capsules, capsules were first transferred to gelatin or BSA coated petri-dishes filled with ASW. Fine forceps were used to carefully tear the capsule apart without damaging the embryos until a large enough opening is created for the embryos to be released into the dish. Debris from the capsules or damaged embryos were promptly removed from the dish during the process.

Embryo fixation

Embryos were decapsulated when they reached the desired stages. Decapsulated embryos were transferred to fixative containing tubes with a coated pipette tip. Eppendorf lo-Bind 2mL tubes (022431048, 022431102, eppendorf) were used to contain up to 60 embryos per tube. Two fixatives were prepared: the first with 4% formaldehyde in 2x PBST, and the second with 4%

formaldehyde in 1x PBST (0.1% final tween concentration for both) plus RNase inhibitor (M0314S, NEB) at 1:1000 to 1:5000 dilution. Embryos were first transferred to the fixative with 2x PBST (to reduce shock of transitioning from the high salinity of sea water to low salinity of 1x PBST). After 1-5 minutes, the initial fixative was removed from the tube and replaced with the second fixative (4% formaldehyde in 1x PBST with RNase inhibitor). Embryos were fixed at 4°C overnight (14-20 hrs). The next day, embryos were washed 3 times (5 minutes each) in PBST, then dehydrated over a few washes to replace PBST with 100% methanol (10 minutes with 50% PBST 50% methanol, followed by 3 times 5 minutes each with 100% methanol). Embryos were stored in methanol at -20°C until used for MERFISH or immunostaining.

PolyA-FISH-coupled immunostaining and imaging

Fixed and dehydrated embryos were retrieved from -20°C and equilibrated to room temperature. Embryos were rehydrated by first washing in a 50% methanol and 50% PBST solution for 10 minutes, followed by 3 washes in PBST. An antigen retrieval step was performed by incubating embryos in 150mM Tris-HCl buffer (pH 9) at 70°C for 12 minutes. Embryos were washed briefly in PBST again before incubation in the blocking solution (3% BSA in PBST, with 1:1000 - 1:5000 RNase inhibitor). Blocking was performed either at 4°C overnight or at room temperature for 2-3 hours. Primary antibody solution was prepared by diluting primary antibodies and polyT-488 probe (for polyA-FISH, /5Alex488N/TTTTTTT+TTT+TT+TT+TTT+TTTTTTTT, IDT) in the blocking solution. PolyT-488 probe stock at 100uM was diluted at 1:2000. Antibodies for α-tubulin (MCA77G, bio-rad), pan-actin (AAN02, cytoskeleton), β-catenin (C2206, sigma), γ-tubulin (T3559, sigma) were diluted at 1:400. Primary antibody solution was passed through a 0.22um syringe filter (09-720-3, Fisher) before use. Embryos were incubated in the primary antibody at 4°C at least overnight. Next, embryos were washed with PBST 6 times, 15 minutes each (or 5 times, 20 minutes each). Secondary antibody solution was prepared by diluting polyT-488 probe and secondary antibodies in PBST with 1% BSA and 1:5000 RNase inhibitor. Cross-absorbed secondary antibodies (A21247, A21037, Invitrogen; 615-165-214, Jackson Immuno Research Labs) were used at 1:500 dilution. Secondary antibody solution was passed through a syringe filter before use. Embryos were incubated in the secondary antibody solution either at 4°C overnight or at room temperature for 3-5 hours. Embryos were then washed with PBST 6 times, 15 minutes each (or 5 times, 20 minutes each), and stained with DAPI (MBD0015, Sigma) at 1:3000 dilution for 2 hours at room temperature or overnight at 4°C. Embryos were briefly rinsed in PBST before mounted for imaging. To achieve optical clarity, embryos were dehydrated in methanol or ethanol, then soaked in the refractive index matching media BBBA (1:1 benzyl benzoate : benzyl alcohol mixture). Embryos were mounted on a coverslip with a shallow chamber made by cutting a hole in the center of two layers of electric tape (which matches roughly the thickness of the embryos). Excess BBBA were added with the embryos to fill the hole, and another coverslip was gently placed on top to seal the chamber without introducing bubbles. The sample was then imaged on the Leica Stellaris 5 Confocal microscope using 20x objective and with default built-in Lightening processing.

Transcript annotation

*C. atrasolea* transcripts were named after the best blast hits of their predicted proteins against the uniprot database. Proteins encoded by each transcript were predicted using

Transdecoder (https://github.com/sghignone/TransDecoder). Reference proteome data from diverse organisms (uniprot_sprot.fasta) was downloaded from https://www.uniprot.org/help/downloads in fasta format under the UniProtKB section Reviewed (Swiss-Prot) resources (*71*). A blastable database was made from the downloaded proteome data using the *makeblastdb* command from the BLAST command line application (*72*). Blasting was done using the *blastp* command with the following options: *sta -evalue 1e-5 -max_target_seqs 5 -outfmt 6*. The top hit by p-value was used for annotating each transcript. No blast hits were found for two transcripts using this approach. For these, we performed blastx using NCBI webtool. Both transcripts returned gastropod proteins not present in the uniprotKB as their best hits. We named these transcripts after their best blastx hits as 'hypothetical protein' and 'uncharacterized protein'.

#### Genomic and transcriptomic data for probe design

The *de novo* assembled reference transcriptome is available from GenBank under the accession GKTO00000000. A genomic DNA sequencing dataset by PacBio is deposited in the NCBI Sequence Read Archive (SRA) under accession number PRJNA1529796. A bulk RNA sequencing dataset containing 12 early developmental stages is deposited in the NCBI Gene Expression Omnibus (GEO) repository with accession number GSE347490.

#### Candidate gene selection for MERFISH

Gene panels for spatial transcriptomic assays were commonly selected from cell type markers based on single-cell RNA-seq data. Genes could be additionally picked based on their functions. However, no single-cell RNA-seq data was available for any spiralian embryos at cleavage stages. Additionally, annotations of *C. atrasolea* transcripts based on their single best match in annotated species failed to resolve the functions of many genes due to their low homology to reference genomes/transcriptomes and lack of functional studies in closely related species, which is a common challenge in non-model organisms. Despite these challenges, we selected candidate *C. atrasolea* genes for MERFISH using three approaches. First, a bulk RNA-seq dataset containing samples from early developmental stages (2-cells to 53-cells) were used to define 7 gene sets with distinct developmental dynamics. Ten genes from each set with expression levels below a threshold (<500 TPM) were selected as MERFISH candidates. Second, a list of manually curated keywords for genes or functions classically involved in developmental and cellular processes were searched against *C. atrasolea* transcript annotations. Transcript annotations were generated using InterproScan, which summarized predicted protein family membership, functional pathways, and the presence of functional domains and sites for each transcript (*73*). Over 30 keywords were searched, including developmental signaling pathway components, such as 'bmp', 'notch', 'fgf', transcriptional regulators, such as 'sox', 'smad', 'homeobox', and cell cycle regulators, such as 'cyclin'. Transcripts whose annotations included the searched keywords and were expressed in at least one stage (TPM >5) were picked as MERFISH candidates. Third, protein sequences of 50 selected developmental regulators, housekeeping genes, cell cycle regulators, and genes with known subcellular locations were curated from organisms with relevant functional studies. Some sequences were collected from model organisms such as mouse (e.g. *actin*), fish (e.g. *ddx6*), and fly (e.g. *cen*), while others were picked from spiralian species such as Crepidula fornicata (e.g. *dpp*), Owenia fusiformis (e.g. *delta*), and Ilyanassa obsoleta (e.g. *iotis11*). Their orthologous proteins in *C. atrasolea* were identified through blast search. Transcripts corresponding to the best hits were selected as MERFISH candidates. The three approaches returned a union of 452 candidates. Python scripts for candidate selection are deposited in Dryad (DOI: 10.5061/dryad.dz08kpsdn). Importantly, since no *C. atrasolea* genome was available, candidates were picked from a *de novo* assembled transcriptome with no reliable gene models. Therefore, the 452 candidate transcripts included both transcripts from distinct genes and isoforms of the same genes. The redundancies between gene isoforms were resolved during probe design. The final set of genes assayed in MERFISH are included in table S1.

<u>MERFISH primary probe design</u>

Our MERFISH assay employed a 3-layer probe strategy, with primary probes directly hybridizing to the target transcripts, gene-specific adaptor probes that bind the primary probes, and readout probes tagged with fluorophores and bind the adaptor probes. Primary probes targeting candidate transcripts were 140nt in length. Each contained a 40nt transcript targeting sequence, three repeats of a 20nt adaptor sequence, and a 20nt primer sequence at each end of the probe for PCR amplification. For each target transcript, 20-60 probes were designed with non-overlapping transcript targeting sequences and a gene-specific adaptor sequence. All probes shared the same primer sequences. Adaptors and primers were picked from a database of existing adaptor and primer sequences from the Bintu lab. Available adaptor and primer sequences were blasted against the *C. atrasolea* RNA- and genomic sequencing data to select the ones with low chance of targeting the endogenous transcriptome or genome. The transcript targeting sequences of the primary probes were designed using a custom pipeline that entailed four major steps. First, coding sequence (CDS) of transcripts were defined and isoforms were predicted for each candidate transcript based on their CDS similarities. Second, possible 40-mer probe sequences were generated from the CDS and the region within 2000 nt of the CDS. Third, the probes were filtered by thresholding on their melting temperature, GC content, as well as their off-target rates in 1) the transcriptome (excluding predicted isoforms), 2) in a genomic sequencing dataset, and 3) in a list of pre-defined repetitive sequences. Finally, redundant probesets (due to isoforms or genes with similar sequences) were removed. This was done by blasting the probeset for each candidate transcript against the transcriptome, grouping probesets with similar hit profiles, and keeping one representative probeset per group. This resulted in the final probesets for 216 unique transcripts. Two non-overlapping probesets were designed for some transcripts for quality control. Each probeset contained 7 to 60 probes, with a median 46 probes per set. In total, the primary probe library contained 11967 probes. Python scripts for probe design are deposited in Dryad (DOI: 10.5061/dryad.dz08kpsdn). Four additional primary probes were designed separately, each targeting the ribosomal rRNA 18S, the mitochondrial rRNA 16S, the spliceosome snRNA U6, or polyA. Only one probe targeting a highly conserved 40nt sequence was designed for 18S, 16S, and U6. The probe for polyA contained a 21nt repeat of thymine flanked with a sequence that hybridizes directly to a polyA readout probe without the need of an intermediate adaptor probe.

<u>MERFISH primary probe synthesis</u>

Probes were prepared from an oligonucleotide pool containing the designed primary probe sequences, which are the reverse complement of the final probes (Twist Biosciences). We largely followed previously published protocols for probe synthesis (*29, 74*), but used a 5'-acrydite-modified primer to incorporate acrydite into the final probes. Briefly, qPCR was first performed on the oligonucleotide pool with a 5'-acrydite-modified reverse primer and a forward primer extended with a 5' T7 promoter sequence. The resulting qPCR product, with acrydite on the reverse strand and T7 promoter on the forward, was cleaned up with the DNA clean & concentrator-5 kit (D4003, Zymo Research). Next, *in vitro* transcription was performed with a T7 high yield RNA synthesis kit (E2040S NEB) to amplify the forward stand sequences as RNAs. This was followed by reverse transcription using the Maxima reverse transcriptase (EP0751, Thermo Fisher) with the 5'-acrydite-modified reverse primer to generate acrydite-modified reverse DNA strand. Alkaline hydrolysis (1:1 of 1M NaOH and 0.5M EDTA) was then performed to remove the RNA templates, and the acrydite-modified DNA probes were purified using the oligo clean & concentrator kit (D4060, Zymo Research). Purified probes were quality checked on a Tapestation D1000 DNA ScreenTape. Probe concentration was measured with a nanodrop. Finally, a SpeedVac was used to concentrate the probes to a final concentration of 3-4ug/uL. Probes were stored at -20°C.

MERFISH primary probe hybridization

Fixed and dehydrated embryos were retrieved from -20°C and equilibrated to room temperature. Embryos were rehydrated by first washing in a 50% methanol and 50% PBST solution for 10min, followed by 3 washes in PBST with RNase inhibitor (1:1000-1:5000 dilution, M0314S, NEB). Embryos were further permeated by a 10 min wash in PBS solution with 0.25% triton X-100 before a 5 min rinsing in 2x SSC. Rinsed embryos were washed twice (10 min each) in the 40% hybridization wash buffer (2xSSC, 1:5000 RNase inhibitor, 0.1% tween, 40% formamide) before incubation in probe hybridization solution. Probe hybridization solution was made by mixing 27 uL probe solution and 3 uL RNase inhibitor with 120 uL hybridization buffer. To make 800 uL hybridization buffer, combine 500uL formamide (17899, Thermo Fisher), 140uL 50% w/v dextran sulfate (S4030, Sigma), 100uL 20x SSC, and 60 uL nuclease free water, mix well before use. The probe solution contained 25 uL synthesized primary probe library at \~4.5 ug/uL concentration (targeting \~6.5 ng/uL per probe concentration in the final probe hybridization solution), plus the four probes targeting polyA (total mRNA), 18S (rRNA), 16S (mitochondrial rRNA), and U6 (spliceosome), with 0.5 uL of each at 100 uM stock concentration. Up to 40 Embryos were incubated in 150 uL probe hybridization solution per 2 mL eppendorf lo-Bind tubes at 47°C overnight (18-24 hrs). The next day, embryos were washed extensively with the 40% hybridization wash buffer with 2 brief washes at 47°C, followed by 6 additional washes totaling 1 hour. Washed embryos were then transferred to a BSA coated petri-dish filled with 2x SSC (with 1:5000 RNase inhibitor) and examined under a dissection microscope. Embryos that looked intact and free of debris stuck on the surface were collected into a fresh 2 mL lo-Bind tube filled with 2x SSC (with 1:5000 RNase inhibitor). Embryos were generally embedded in acrylamide gel within 30 min afterwards.

Probe-hybridized embryo embedding

After primary probe hybridization, embryos were embedded on silanized coverslips in a thin layer of acrylamide gel. To prepare silanized coverslips, No.1.5 coverslips were first cleaned in a mixture of methanol and HCl (1:1 v:v 100% methanol : 37% HCl) by a 30 min incubation at room temperature, then rinsed 3 times with Milli-Q water, followed by 3 rinses in 100% ethanol. Excess ethanol was blown off and the clean coverslips were dried at 70°C for at least 25 min. A

1% silane coupling solution was made by mixing 10 uL TMSPMA (M0725, TCI), 1 mL ethanol, and 100 uL glacial acetic acid. Cleaned coverslips were silanized by immersion in the 1% silane coupling solution at room temperature for 30 min, then rinsed 3 times in ethanol. Excess ethanol was blown off and coverslips were incubated in a dry oven at > 70°C for at least 30 min. Silanized coverslips were kept in the oven until use and were used within 2 days. Acrylamid gel buffer was made by mixing 1.2 mL 1 M Tris buffer (pH8), 1.2 mL 5 M NaCl solution, 4 uL RNase inhibitor (M0314S, NEB), and 2 mL 40% acrylamide (1:19 crosslinking), then adding water to make up a total volume of 20 mL. The gel buffer was degassed in a vacuum chamber to remove excess oxygen prior to use. Embryos were rinsed in the degassed gel buffer twice. 10 mL of the remaining degassed gel buffer was mixed with 50 uL TEMED and 5 uL freshly made 10% w/v APS (A3678, Sigma) solution in water to initiate the gel polymerization process. Embryos were washed with the initiated gel buffer twice quickly and transferred with excess initiated gel buffer to the center of a silanized coverslip using BSA coated pipette tips. A No. 0 coverslip smaller than the silanized cover and coated with Gel Slick solution (50640, Lonza) was gently placed on top to sandwich the embryos and initiated gel buffer without crushing the embryos. The thickness of the gel was controlled by the volume of the gel-embryo mixture transferred to the silanized cover. Since embryos are approximately 300 um along the animal-vegetal axis, we aimed for a 325 um space between the Gel Slick coated and the silanized coverslips. For instance, when a 18mm square Gel Slick coated coverslip was used, 105 uL embryo-gel mixture was transferred. The sandwiched setup was then covered with a petri-dish lid to reduce drying and let sit for 3-6 hrs at room temperature for acrylamide polymerization. The sample was then immersed in 2x SSC in a petri-dish for at least 5 min before the Gel Slick coated cover was carefully lifted and peeled off.

Sequential immunostaining of embedded embryos

Immunostaining was performed on three batches of the MERFISH probe-hybridized, gel-embedded embryos prior to enzymatic sample clearing. Immunostaining followed the protocol described in “PolyA-FISH-coupled immunostaining”, with extended primary and secondary antibody incubation time (3-5 days at 4°C) to allow antibody penetration through the gel-embedded embryos. After DAPI labeling, samples were imaged in PBST on a ZEISS LSM700 confocal microscope. No index matching medium was used for optical clearing due to the formation of opaque residuals or dramatic expansion of the acrylamide gel caused by the medium tested, including BBBA (1:1 benzyl benzoate : benzyl alcohol mixture) and CUBIC-R+(N) (T398325ML, TCI). To multiplex, samples were co-stained with two primary antibodies raised in different hosts at a time, followed by imaging, antibody stripping, and staining of additional antibodies. Antibody stripping was performed with a 2-Mercaptoethanol/SDS (2-ME/SDS) buffer similar as previously published (*75*). The 2-ME/SDS buffer was freshly made before use by combining 1 mL 10% w/v SDS in water, 250 uL 1M Tris-HCL (pH 6.8), 3.7 mL distilled water, and 30 uL 2-Mercaptoethanol. Sample was immersed in the 2-ME/SDS buffer and incubated at 50°C for 2.5 hrs with gentle shaking. Afterwards, the sample was washed in PBSTx (PBS with 1% triton X-100) for 2 hrs at 47°C, followed by additional washes in PBST (PBS with 0.1% tween) at room temperature for 1 hr. The sample was then subjected to a new round of immunostaining from the blocking step. Up to three rounds of immunostaining with 6 primary antibodies in total was performed on the same sample.

Enzymatic clearing of embedded samples

Embedded embryos were cleared with a sample digestion solution made by combining 7.2 mL 2x SSC, 2.5 mL 10% w/v SDS, 100 uL Proteinase K, and 200 uL triton X-100. Samples were immersed in the digestion solution at 60°C for 1 hr, then at 47°C for 16-20 hrs with fresh digestion solution. Samples were then washed in 2xSSC at 47°C 3 times, 20 min each, then kept in fresh 2xSSC at room temperature until mounting for on-stage sequential hybridization and imaging. When the sample could not be mounted within 1 day, it was stored at 4°C until ready to be mounted.

Fluidics-coupled microscope setup for MERFISH

A microscope system was built with an ASI microscope body. Fine translation of the samples will be performed with XYZ stages (ASI). A high numeric aperture (NA = 1.05), long working distance, silicon-oil based 25X objective (CFI Plan Apochromat Lambda S 25XC Sil for biological microscopes, Nikon) was installed for 3D MERFISH imaging. A custom tube 300-mm tube lens was installed to increase the resolution of single molecule localization with an effective magnification of 37.5X. Illumination was provided by a Lumencor CELESTA light engine (wavelengths: 405, 477, 546, 638, 749 nm) with a penta-bandpass dichroic (IDEX, FF421/491/567/659/776-Di01) and filter (IDEX, FF01–441/511/593/684/817). The laser output emerging out of a 400um square fiber was reimaged onto the sample space through the 25X objective coupled with the 300-mm tube lens to ensure uniform epifluorescence imaging across the area captured by the camera. A high-throughput scientific CMOS camera (Kinetix-model from Photometrics) was used for image acquisition. Custom-built autofocus systems employing an IR illumination (Thorlabs LP980-SF15) and a CMOS camera (Thorlabs, uc480) were used to maintain a consistent z-position reference over the many rounds of imaging (*76*). The FCS2 chamber system (Bioptechs) was used to hold the sample during on-stage hybridization and imaging. The chamber was connected to a custom-built, automated fluidics system that consists of two 24-way valves (Vici) and a syringe pump (Hamilton). Each valve was connected to a distinct buffer, with dedicated lines for adaptor probe mixtures, fluorescent readout probes, stripping, washing and imaging buffers. Buffers were delivered to the sample chamber, and waste was collected downstream in a waste bottle. All hardware components were synchronized and controlled using a National Instruments Data Acquisition card (NI PCIe-6353) and custom software based on https://github.com/ZhuangLab/storm-control (*77*). Network attached storage devices (Synology) with >200GB capacity each were used to store and access data acquired by the microscope within a 10GB ethernet network framework connecting the microscopes, the storage devices and the analysis computers.

MERFISH codebook design

To determine which genes will be multiplexed in each round of MERFISH imaging, a codebook was designed. To avoid crowding of single molecule signals, genes with very high expression levels in the developmental bulk RNA-seq data were excluded from multiplexing. Specifically, transcripts whose expression was over 100 TPM at any stage over 70 TPM in two or more stages were excluded. Some targets were then manually added or removed based on smFISH results in prior assays. In total, 192 genes were included in the codebook design, which were equally distributed among three imaging channels (750, cy5, and cy3). We planned 14 rounds of multiplexed hybridization and imaging, with each gene imaged in exactly 4 rounds. For each channel, each gene was assigned a unique 14-bit binary barcode with 4 ones to define which 4 rounds of hybridization it will be targeted. The barcodes were generated such that the Hamming distance between any pair is at least 4. Barcode assignments to genes were optimized using a Metropolis-Hastings-based algorithm to reduce the combined expression level (according to bulk RNA-seq) of genes targeted in the same imaging round. The three channel-specific 14-bit matrices were interleaved to form a 42-bit codebook. Eighty 'blank barcodes' were added to the codebook to control for false positives in decoding. Similar to the gene barcodes, the blank barcodes are 14-bit binaries with 4 ones, each with Hamming distances of at least 4 from every gene barcode. Script for codebook design is deposited in Dryad (DOI: 10.5061/dryad.dz08kpsdn).

MERFISH adaptor and readout probe preparation

Adaptor probes and readout probes for MERFISH were purchased from IDT and stored at 100uM concentration. Adaptor probes for genes targeted in the same hybridization round across the three channels were pooled according to the codebook. Pooling of adaptor probes was done using a custom pipetting robot built by combining a CNC Router (CNC-4040-Basic, FoxAlien) with a programmable syringe pump (Hamilton). This robot draws gene-specific adaptor probes from 384 plates and mixes them in 2mL Lo-Bind tubes corresponding to MERFISH hybridization rounds. Code for operating this custom pipetting robot is available from github (https://github.com/BogdanBintu/PipettingRobot). Genes not included in the multiplexed imaging rounds were imaged one per imaging channel per hybridization round. Adaptor probes for individual genes imaged in the same round across color channels were mixed manually. Each adaptor probe contains sequences reverse complementary to one of the three readout probes. Each readout probe is tagged with one of the following three fluorophores: Alexa-750, Cy5, and Cy3. An additional readout probe tagged with Alexa-488 specifically targeting the polyA primary probe was used in each round of hybridization (except for experimental batch 1). During on stage hybridization steps, pooled adaptor probes and readout probes were diluted in a hybridization buffer (0.1% Tween-20, 35% formamide, 2xSSC in milli-Q water) to make working solutions. Adaptor probe solution contains approximately 50nM per adaptor probe, while readout probe solution contains 50nM of each readout probe (1:2000 dilution from the stock).

On-stage sequential hybridization and imaging of MERFISH

The embryo-embedded coverslip was mounted in a FCS2 chamber (Bioptechs) for on-stage hybridization and imaging. A 1 mm thick hexagonal gasket (Bioptechs) was used between the sample coverslip and the microaqueduct slide. The chamber was connected to a fluidics system (see "Fluidics-coupled microscope setup for MERFISH"). Multiple rounds of on-stage hybridization and imaging was done by repeating the following steps: 1. Pre-hybridization, 2. Adaptor probe hybridization, 3. Adaptor probe wash, 4. Readout probe hybridization, 5. Readout probe wash, 6. Imaging, and 7. Probe stripping. Buffer exchanges by the fluidics system were under control of the custom software, called Kilroy (*77*). 1-2 mL buffer was pulled in each exchange. Pre-hybridization was done by incubation in the hybridization wash buffer (30% formamide, 1% tween, 2x SSC in milli-Q water) for 5 min. Adaptor probe hybridization was done by incubation in the adaptor probe solution for 65 min. Readout probe hybridization was done by incubation in the readout probe solution for 35 min. Each probe wash step was done by 10 exchanges of the hybridization wash buffer over 2.5 hrs. At the beginning of the imaging step, the sample was first washed in 2x SSC for 1 hr, followed by equilibration in the imaging buffer (2x SSC, 10% glucose, 50 mM Tris-HCl, 1.6 mM Trolox, 50 uM trolox quinone, 0.3 mg/mL glucose oxidase, 40 ug/mL catalase, and 2 ug/mL DAPI) for 30 min. Stripping was done by 3 exchanges of probe stripping buffer (80% formamide, 0.8x SSC, 1% tween in milli-Q water) over 2 hrs followed by washing with 2x SSC for 5 min. During the imaging step, each embryo was imaged in one field of view up to 400 um deep with 0.5 um step size in Z. MERFISH readout probes carrying cy3, cy5, and Alexa-750 fluorophores were excited by 546 nm, 638 nm, and 749 nm lasers, respectively; polyA readout probe conjugated with Alexa-488 and DAPI were excited by 477 nm and 405 nm lasers, respectively. Five batches of MERFISH experiments were done. Experimental batch 1 entailed 16 rounds of multiplexed hybridization, while the other experiments performed 14 rounds of multiplexed hybridization with an updated codebook. In each experiment, up to 10 rounds of smFISH with one gene per channel was performed after the multiplexed rounds. These single-gene smFISH included some genes that were not imaged in the multiplexed rounds due to high expression levels (indicated by bulk RNA-seq). Some genes that were imaged in the multiplexed rounds were imaged again during the single-gene rounds for benchmarking the decoded results, and some were imaged with alternative probesets for quality control of the probe design. A final round of hybridization was done for the rRNA 18S (Alexa-750), the mitochondrial rRNA 16S (cy5), and the spliceosome snRNA U6 (cy3).

Image deconvolution

The raw MERFISH images are deconvolved using empirical PSFs and the python sdeconv module (https://github.com/sylvainprigent/sdeconv). Experimental PSFs were computed from 28 smFISH images with Alexa-750 conjugated readout probes. Genes were imaged one gene at a time (without MERFISH multiplexing) to favor non-overlapping single-point signals for PSF computation. The images were acquired across 3 experimental batches, 21 embryos spanning 1-cells to organogenesis stages, and 11 genes. Four PSFs were computed over the maximum imaging depth of 400 um: within the first 100 um, between 100-200 um, between 200-300 um, and between 300-400 um. PSFs were computed with a custom script that first identifies isolated point signals by local maximum searching, extracts a volume around each one, precisely aligns their centers using Fourier shifts, then averages the ones from the same Z-range to produce a 3D PSF, and finally cleans the PSF by removing per Z-slice and broad background and normalizes the cleaned PSF such that the final empirical PSF values range between 0 (background) and 1 (peak). Overall, 24635, 18667, 9858, and 23363 point signals were used to produce PSF1 (from 0-100 um), PSF2 (100-200 um), PSF3 (200-300 um), and PSF4 (300-400 um), respectively. Image deconvolution was performed using the sdeconv function *deconv.SRichardsonLucy* with the empirical PSFs. Four overlapping chunks along Z were first extracted from each 3D image array: 0-125 um, 75-225 um, 175-325 um, and 275-400 um. These four image chunks were deconvolved using PSF1, PSF2, PSF3, and PSF4, respectively. The resulting deconvolved image chunks were then stitched back together, with a linear function combining the overlapping regions as the weighted average to achieve a smooth transition (with the weight going from 0 to 1 for one image and from 1 to 0 for the other over the overlapping Z). Scripts used for PSF computation and deconvolution are deposited in Dryad (DOI: 10.5061/dryad.dz08kpsdn).

MERFISH point detection and global drift correction

Point signal detection and drift correction followed previously described pipelines (*28, 78*). Briefly, point signals representing single transcripts were detected from deconvolved images as Gaussian peaks. Decoding transcript identities of the point signals requires the spatial alignment of the same points across hybridization rounds. To correct for the spatial drift of images between hybridization rounds, we first estimated the drift using signals detected in the nuclei (DAPI, 405 channel) images acquired in conjunction with the transcript channels; polyA signals (488 channel) were additionally used when available. DAPI and polyA features corresponding to local minima and maxima intensity were detected with Gaussian peak fitting. Phase cross-correlation was performed on these features between one reference round and each of the other rounds to infer spatial drift. The inferred drifts were subtracted from the point signal coordinates to register points across hybridization rounds. Scripts are deposited in Dryad (DOI: 10.5061/dryad.dz08kpsdn).

MERFISH local distortion correction

In addition to the global drift correction, local distortion correction was performed to address the non-uniform expansion or contraction between hybridization rounds in the thick sample. After applying the global drift correction, each 3D image volume was divided into a regular grid of local subvolumes. Between the reference hybridization round and each other round, the displacement of each local subvolume was estimated through phase cross-correlation of DAPI and polyA features, similar to how global drift was inferred. The centroid of the reference subvolumes and their displacements relative to another hybridization round were used to fit a 3D cubic B-spline transformation. The fitted transformation provided a continuous and smooth displacement function throughout the imaging volume. This function was applied to the coordinates of all point signals to correct for local distortion effects. Correction was assessed by visualizing DAPI and polyA feature alignment with the reference round before and after correction. Scripts are deposited in Dryad (DOI: 10.5061/dryad.dz08kpsdn).

Iterative MERFISH data decoding

Following the initial feature-based local distortion correction, the point signals detected in MERFISH were decoded for their transcript identities and their spatial registration was iteratively refined using confidently decoded transcripts as registration markers. In each iteration, point signals were decoded as described previously (*28, 78*). Briefly, points from different hybridization rounds were grouped into candidate molecules based on their spatial proximity. Candidate molecules were assigned transcript identities by comparing their detected-bit patterns with the four-bit barcodes in the MERFISH codebook. The initial decoding used a relaxed maximum grouping distance of 5 pixels. A decoding quality score incorporating signal brightness, spatial compactness, and agreement with the expected barcode was calculated for each decoded transcript, and transcripts with scores greater than -2 were retained as registration markers. For each non-reference hybridization round, transcripts containing signals in both that round and the reference round were identified. The coordinates of the two signals belonging to the same decoded transcript provided a direct estimate of the local displacement between the hybridization rounds. For every detected point, a local correction was estimated from the 200 nearest displacement measurements. Neighbor contributions were weighted by a Gaussian function of their spatial distance, with greater weight assigned to nearby measurements. The estimated displacement (with a maximum displacement cutoff of 9 pixels) was subtracted from point signals in the corresponding hybridization round, whereas points from the reference round were left unchanged. Transcript identities were decoded again after each correction, and the new decoding was used to estimate the local displacement in the subsequent iteration. Four to seven rounds of correction and decoding were performed. The maximum distance used to group signals into candidate transcripts during decoding was progressively reduced from 5 to 3 pixels over the iterations. Effects of the distortion correction were assessed by comparing the number of molecules decoded, false positive (blank barcode) rate, and signal displacements across iterations. A final round of decoding was performed on the iteratively corrected point signals, with the maximum grouping distance set to 2 pixels.

Label-free RNA community analysis

To characterize the spatial organization of transcripts without relying on segmented cellular or subcellular structures, we grouped individual mRNA molecules into local neighborhoods, termed communities, and clustered these communities by their transcriptional composition into RNA domains. RNA molecule coordinates were converted from voxels to microns (voxel dimensions z = 0.5 um; x, y = 0.177 um). Communities were constructed by nearest-neighbor search with a k-d tree using scipy.spatial.KDTree module. Every 37th molecule was taken as a community center and the 350 nearest molecules to each center were retrieved, excluding neighbors farther than 30 um. At these settings each molecule was contained in a median of 9 communities. For experimental batch 2, which yielded sparser transcript detection, a community size of 300 was used so that community radii remained comparable across batches. The resulting gene by community count matrix was subjected to PCA and a k-nearest-neighbor graph (K = 30) was built for the communities with the top 18 PCs using cosine similarities as edge weights. The community graph was partitioned with the Leiden algorithm using the leidenalg module to produce the community clusters. Each molecule was then assigned the cluster identity of its nearest community center, excluding clusters consisting of a single community. This yields a domain label for every molecule in the embryo. Domains of few molecules (< 15) were pooled into a residual label that was excluded from downstream analysis. Domains were annotated manually according to their spatial patterns to reflect their cellular and subcellular identities. The community analysis was performed on each embryo twice: first to identify and remove fluorescent contaminants (see "Computational removal of fluorescent contaminants"), and then to produce the final RNA domains from the contaminant removed data. Scripts for performing the Label-free RNA community analysis are deposited in Dryad (DOI: 10.5061/dryad.dz08kpsdn).

Computational removal of fluorescent contaminants

MERFISH and single-molecule FISH data sometimes included fluorescent signals from dust, lint, probe aggregate, or other contaminants. Signals from such contaminants often were more intense compared to real single-transcript signals and manifested as large hourglass-shaped patterns along the Z-axis even after image deconvolution. We removed such contaminant signals by applying the community analysis described above, using the domain spatial pattern and domain molecule decoding rate to identify contaminants. For each domain we computed the proportion of blank barcodes, which reflects the false positive rate from decoding, as well as the mean MERFISH decoding score. Domains with the characteristic large hourglass shape in the 3D space and elevated blank barcode proportion or low decoding score were annotated as contaminants. Domains annotated as contaminants had a median blank proportion of 0.014, compared with 0.0041 for retained domains, and median decoding score of -1.61, compared with -1.33 for retained domains. All molecules in annotated contaminant domains were discarded. This community analysis, contaminant annotation and removal pipeline was repeated once more. Points corresponding to the blank barcodes were additionally removed. The remaining molecules constituted the cleaned data used for all downstream analyses and plotting in the paper, including cell type clustering, marker gene identification, and the final RNA domain analysis.

Cell segmentation

To segment cells in the 3D embryo, we used a custom pipeline that combines the output of neural-network based cell segmentation, Cellpose4 (*79*), with a custom transcript-based cell assignment algorithm. We leveraged three modes of data for each embryo assayed in MERFISH, 1) the deconvolved 3D DAPI image capturing the nuclei; 2) A 3D image with the total mRNA generated either using the polyA probe or using a 3D density map of the MERFISH-decoded transcripts, and 3) the 3D localizations of the detected MERFISH transcripts. We first use the 3D DAPI image for segmenting the nuclei of each cell. We normalized the deconvolved DAPI image using local background subtraction and then thresholded the normalized image to determine signal within the nuclei. We used density-based clustering in python (DBSCAN) to isolate the nuclear signal points and used 3D convex hulls to fill in the 3D volumes of each nucleus. To perform cell segmentation, we associate each transcript to the most probable nucleus using the following procedure: 1) We used Cellpose4 to get the estimated gradients of each cell from the total RNA image. 2) We processed the Cellpose-inferred gradients using a Sobel kernel to get a 3D edge probability image. 3) We connected each MERFISH detected transcript to each candidate nucleus through a set of 3D lines called travel lines. 4) For each travel line we computed the edge travel score defined as the sum of the signal covered by the travel line in the 3D edge probability image. This gives a measure of how many cell boundaries were crossed as we connected each transcript to each candidate nucleus. We assigned each transcript to the nucleus that minimizes the edge travel score. Scripts for performing the cell segmentation are deposited in Dryad (DOI: 10.5061/dryad.dz08kpsdn).

Cell lineage annotations

Cells were annotated based on their embryonic positions and morphologies according to spiral cleavage conventions. Assignment of quartet identities was trivial. However, from the 4- to 24-cells stages, embryos exhibit apparent tetraradial symmetry, posing challenges to quadrant identity determination. Assignment of quadrant identities was guided by two clues. First, macromeres from B and D quadrants form large contact with each other at the center of the embryo while A and C are mostly not in contact. This cell-cell contact difference was observed in both the polyA FISH and immunostaining. Second, the D quadrant macromere is slightly more advanced in their cell cycle compared to other quadrants. The cell cycle phase of cells was inferred by their chromosome (by DAPI stain) and cytoskeletal structure (immunostaining) configurations. In most cases, cell lineages could be determined unambiguously. An exception was made for the 2-cells stage embryos, which contain the “AB” and “CD” cells. The two cells exhibit highly similar morphologies and can be distinguished from each other only by the presence of a small transient vegetal protrusion (the polar lobe) specific to the CD cell during mitosis. The polar lobe was only observed in two 2-cells stage embryos in our dataset. We therefore annotated all cells at the 2-cells stage ambiguously as “AB/CD”.

UMAP and leiden clustering of single-cell transcriptomes

Cell segmentation allowed the extraction of single-cell transcriptomes from the MERFISH data. To visualize this high-dimensional data and identify transcriptionally distinct cell clusters, we followed the standard scRNA-seq analysis pipeline to perform UMAP and Leiden clustering. The python module Scanpy was used (*80*). Specifically, the cell-by-gene count data was total-count normalized and log2 transformed (after adding a pseudocount of 1). Because different imaging batches targeted overlapping but non-identical gene panels, combined expression tables were assembled on the 192 genes assayed across all batches. PCA was then performed using all genes, followed by batch correction using the Scanpy function external.pp.bbknn, which implements a batch-balanced K-nearest neighbor algorithm (*81*). This generated a batch-corrected nearest-neighbor graph while leaving the original expression matrix unchanged. UMAP embedding and Leiden clustering were subsequently performed using this corrected neighbor graph.

Marker gene identification

Marker genes for cell lineages were identified from the MERFISH derived single-cell transcriptomes. Analysis was restricted to embryonic cells and polar bodies were excluded. Expression tables were assembled on the union of genes across batches, and genes not measured in a given batch were retained as missing values rather than set to 0. Counts were total-count normalized and log2 transformed (after adding a pseudocount of 1). For each gene, expression was modeled with a robust linear model based on iteratively reweighted least squares with a Huber T estimator (statsmodels) (*82*). The model is formulated as log2(normalized expression) \~ lineage group + experimental batch. Lineage groups of cells were assigned according to their lineage annotations (see Cell lineage annotations). For marker gene identification, cells were grouped either by their quartet identities (1q, 2q, 1Q, 2Q, etc.) or by their quadrant identities (A, B, C, D). The model was fitted using all cells. Fitting each gene returned a coefficient and p-value for every lineage group and for every batch. The lineage group coefficients were then used to compute a group-effect score to evaluate how far the coefficient of a particular lineage group deviated from other lineage groups'. A higher score indicates that the gene is more enriched in the scored lineage group compared to other lineages. Scores can be negative to indicate depletion of a gene in a lineage group. A log2 fold change was additionally calculated per gene per lineage group by taking the log2 ratio between target-group mean count and rest-group mean count (with the addition of pseudocount 1). The log2 fold change was calculated per batch and averaged. Genes were called enriched markers of a quartet lineage group when they satisfied score $\geq 2$ or log2 fold change $\geq 0.8$ together with an adjusted (Benjamini–Hochberg) p value $\leq$ 10e-20. Scripts for these analyses are deposited in Dryad (DOI: 10.5061/dryad.dz08kpsdn).

Fluorescent reporter mRNA preparation

Plasmids were constructed to produce mRNAs encoding fluorescent reporters for live imaging. These plasmids are listed below with their Addgene ID:

1. pCS2-PABP(N)-mNeonGreen, Addgene ID - 262458;
2. pCS2-EMTB-TDmRuby, Addgene ID - 262460;
3. pCS2-H2B-CFP, Addgene ID - 262459;
4. pCS2-membrane-CFP, Addgene ID - 262461;
5. pCS2-LifeAct-CFP, Addgene ID - 262462;

All plasmids contain the Sp6 promoter sequence upstream of the coding sequence of the reporters. mRNAs for injection were made from linearized plasmids using the mMESSAGE mMACHINE SP6 Transcription Kit (AM1340, Invitrogen). We modified the manufacturer's protocol and performed the *in vitro* transcription overnight at 30°C. The resulting mRNAs were polyadenylated using the Poly(A) Tailing Kit (AM1350, invitrogen), then cleaned using the RNA Clean & Concentrator-25 kit (R1017, Zymo) and eluted in nuclease-free water.

Embryo injection and live imaging

The injection mix was prepared to contain 1:5 dilution of 0.5% phenol red solution (P0290, Sigma), as well as mRNAs of interest at their working concentrations (EMTB-TDmRuby: 75ng/uL; PABPN-mNeonGreen: 50ng/uL; H2B-CFP: 50ng/uL; membrane-CFP: 75ng/uL; LifeAct-CFP: 75ng/uL). Injection mixes were made within 3 hrs of injection and maintained on ice until use. Embryos at 1-cell stage were released from their egg cases (see "Embryo decapsulation") and transferred to a petri dish coated with 2% agarose gel with 300um wide and 300um deep lanes to hold the embryos during injection. Glass capillary tubes (TW100F-4, WPI) were pulled into injection needles using Sutter Instrument P-1000 micropipette puller with the following program: Heat = 550, Pull = 80, Velocity =80, Delay = 90, Pressure = 200, Ramp = 538. The injection mix was loaded into the injection needle and approximately 0.25nL was injected into each embryo using the Narishige IM-400 Electric Microinjector. Injected embryos were raised at room temperature until they reached 2-4 cells stage. Meanwhile, a glass bottom MatTek dish (P35G-1.5-20-C, MATTEK) was first coated with BSA or gelatin (see "Coating of dishes and pipette tips"), then coated with 1% low-melt agarose gel with microwells stamped by a 3D printed stamp to hold the embryos in place. The dish was filled with filtered ASW and injected embryos were transferred to the microwells and manually oriented using an eyelash tool. Live imaging was performed on a Leica SP8 confocal microscope in the lightning mode with the dish held in a temperature controlled chamber set at 27°C. Movies were acquired at 7-minute intervals. Up to 3 embryos were imaged per experiment.

Pharmacological treatment assay

Pharmacological inhibition of microtubule polymerization and actin polymerization was done using nocodazole (487928, sigma) and cytochalasin D (5047760001, sigma), respectively. Nocodazole treatment was performed at 20ug/mL final concentration, while cytochalasin D treatment was performed at 12.5ug/mL final concentration. Both drugs were stored as high concentration stocks in DMSO per manufacturer's instructions. Cytochalasin D stock solution was stored at -80°C while Nocodazole stock solution was stored at -20°C in small aliquots to avoid repeated freeze-thaw. Stocks were diluted in filtered (0.22um) ASW to 2X of the final working concentration 1-2 hours prior to treatment. Control media were made by diluting DMSO at the same v/v concentration as the drugs. The 2X drug and control media were filtered through a 0.22um syringe filter (09-720-3, Fisher) and kept at 27°C until use. Embryos injected with fluorescent reporters (e.g. for H2B, membrane, and microtubule) were raised to 4-8 cells stage before transferred to a custom made imaging dish, which allowed live imaging of control and drug-treated embryos in the same setup. The imaging dish was constructed from a glass bottom MatTek dish (P35G-1.5-20-C, MATTEK) by adding a vertical glass divider (cut out from a coverslip) to separate the two halves of the dish. Sylgard 184 elastomer was applied to seal any gap at the bottom and the sides between the divider and the dish to ensure that no liquid exchange occurs between the two sides of the dish. The elastomer was cured at 60°C for 24 hours. The resulting imaging dish was first rinsed in ethanol then cleaned with distilled water. The dish was coated with BSA (see "Coating of dishes and pipette tips"), then coated with 1% low-melt agarose gel with microwells stamped by a 3D printed stamp to hold the embryos in place. Embryos were transferred to each side of the imaging chamber filled with filtered ASW. Live imaging was performed on a Leica SP8 confocal microscope in the lightning mode with the dish held in a temperature controlled chamber set at 27°C. Movies were acquired at 15-minute or 16-minute time intervals. Three to six embryos were imaged from each side of the dish per experiment. Embryos were first imaged in ASW for up to 1 hour, then the dish was carefully taken off the stage. Half of the ASW were removed from each side of the dish and replaced with the 2X DMSO control media on one side and the 2X drug media on the other. The embryos were kept immersed in liquid during the process, and all pipetting steps were done gently to avoid dislocating the embryos. The dish was then placed back on stage to continue the imaging after adjustments were made to correct for small global positional shifts.

Affinity-competition model formulation

We modeled transcript abundance at mRNA aggregates after the well-established multi-component adsorption framework (*45, 46*). Under this framework, at equilibrium, the fractional occupancy of transcripts from the gene g in the aggregation site can be expressed as

$$F_{g,aggr} = \frac{K_g \cdot F_{g,cyto}}{1+\sum_i K_i \cdot F_{i,cyto}} \tag{1}$$

where Ki is the affinity constants of gene i (formally defined as the ratio between the association and dissociation constants), and Fi,cyto is the concentration of gene i in the cytoplasm. The relative occupancy between two genes therefore follows Equation 2.

$$\frac{F_{g1,aggr}}{F_{g2,aggr}} = \frac{K_{g1} \cdot F_{g1,cyto}}{K_{g2} \cdot F_{g2,cyto}} \tag{2}$$

Assuming the aggregation site is saturated at equilibrium, the relative fractional occupancy between genes will remain unchanged, while the sum of the fractional occupancy at the aggregation site over all genes will be 1 (Eq. 3).

$$\sum_i F_{i,aggr} = \sum_i F_{g,aggr} \cdot \frac{K_i \cdot F_{i,cyto}}{K_g \cdot F_{g,cyto}} = \frac{F_{g,aggr}}{K_g \cdot F_{g,cyto}} \sum_i K_i \cdot F_{i,cyto} = 1 \tag{3}$$

Rearranging Equation 3 will derive the formula for the affinity constant K for gene g (Eq. 4).

$$K_g = \frac{F_{g,aggr}}{F_{g,cyto}} \cdot \sum_i K_i \cdot F_{i,cyto} \tag{4}$$

In any given cell, $\sum_i K_i \cdot F_{i,cyto}$ is a constant independent of g. Therefore, the affinity constant K of a gene is proportional to the ratio between the gene's fractional occupancy in the aggregate over its concentration in the cytoplasm. For convenience, we drop the constant term and redefine Kg as the relative affinity coefficient such that

$$K_g = \frac{F_{g,aggr}}{F_{g,cyto}} \tag{5}$$

Due to aggregation site saturation, the fractional occupancy of g in aggregate is equivalent to its fractional abundance, which can be calculated as

$$F_{g,aggr} = \frac{C_{g,aggr}}{\sum_i C_{i,aggr}} \tag{6}$$

where Ci,aggr is the transcript count of gene i in the aggregate. Similarly, the concentration of gene g in the cytoplasm can be calculated as

$$F_{g,cyto} = \frac{C_{g,cyto}}{\sum_i C_{i,cyto}} \tag{7}$$

where Ci,cyto is the transcript count of gene i in the cytoplasm.

<u>Affinity-coefficient calculation</u>

The relative affinity coefficients (Kg) of genes measured in MERFISH were calculated according to equations 5-7. Aggregation and cytoplasmic domains were defined based on the label-free RNA domain analysis. Domains corresponding to subcellular aggregates and cytoplasm were manually annotated according to their spatial patterns. Cell quartet identities were additionally annotated for each domain. Each set of Kg was calculated using domains with the same quartet identity. For instance, a set of Kg values were calculated from 1Q cells in an embryo that contained both 1Q subcellular aggregate domain and 1Q cytoplasm domain. Fg,aggr was calculated using transcripts in the 1Q subcellular aggregate domain, while Fg,cyto was calculated using transcripts in the 1Q cytoplasmic domain, with transcripts within 5 um distance to the subcellular aggregate domain excluded. Kg was calculated only if g has at least 100 transcript counts in both subcellular and cytoplasmic domains. Using this approach, a set of Ks were calculated from each embryo that contained a macromere cell quartet (Q, 1Q, 2Q, or 3Q) with both subcellular aggregate and cytoplasmic domains. K values calculated from the same quartets were averaged per gene to obtain the quartet specific affinity coefficients. These quartet specific affinity coefficients were then averaged (over Q, 1Q, 2Q, and 3Q) to produce the final affinity coefficients shown in Figure 6D.

<u>Subcellular aggregate composition prediction by affinity-competition model</u>

To predict subcellular aggregate composition in a specific cell quartet, a set of relative affinity coefficients (Kg), as well as fractional abundance of genes in the cytoplasm domain (Fg,cyto) are needed. Kg was calculated as described in "Affinity-coefficient calculation", while Fg,cyto was calculated from the RNA domains annotated as the cytoplasm of the target cell quartet. When multiple embryos contain cytoplasm domains of the target cell quartet, fractional abundances were calculated for each embryo then averaged. As in "Affinity-coefficient calculation”, transcripts in the cytoplasm domains within 5 um from the aggregate domains were excluded from Fg,cyto calculation. To predict the fractional abundance of genes in the aggregate domain (Fg,aggr), we use the following formula (Eq. 8) derived from Equation 5 and satisfying the condition that the total fractional abundance summed over all genes equals 1:

$$\hat{F}_{g,aggr} = \frac{K_g \cdot F_{g,cyto}}{\sum_i K_i \cdot F_{i,cyto}} \tag{8}$$

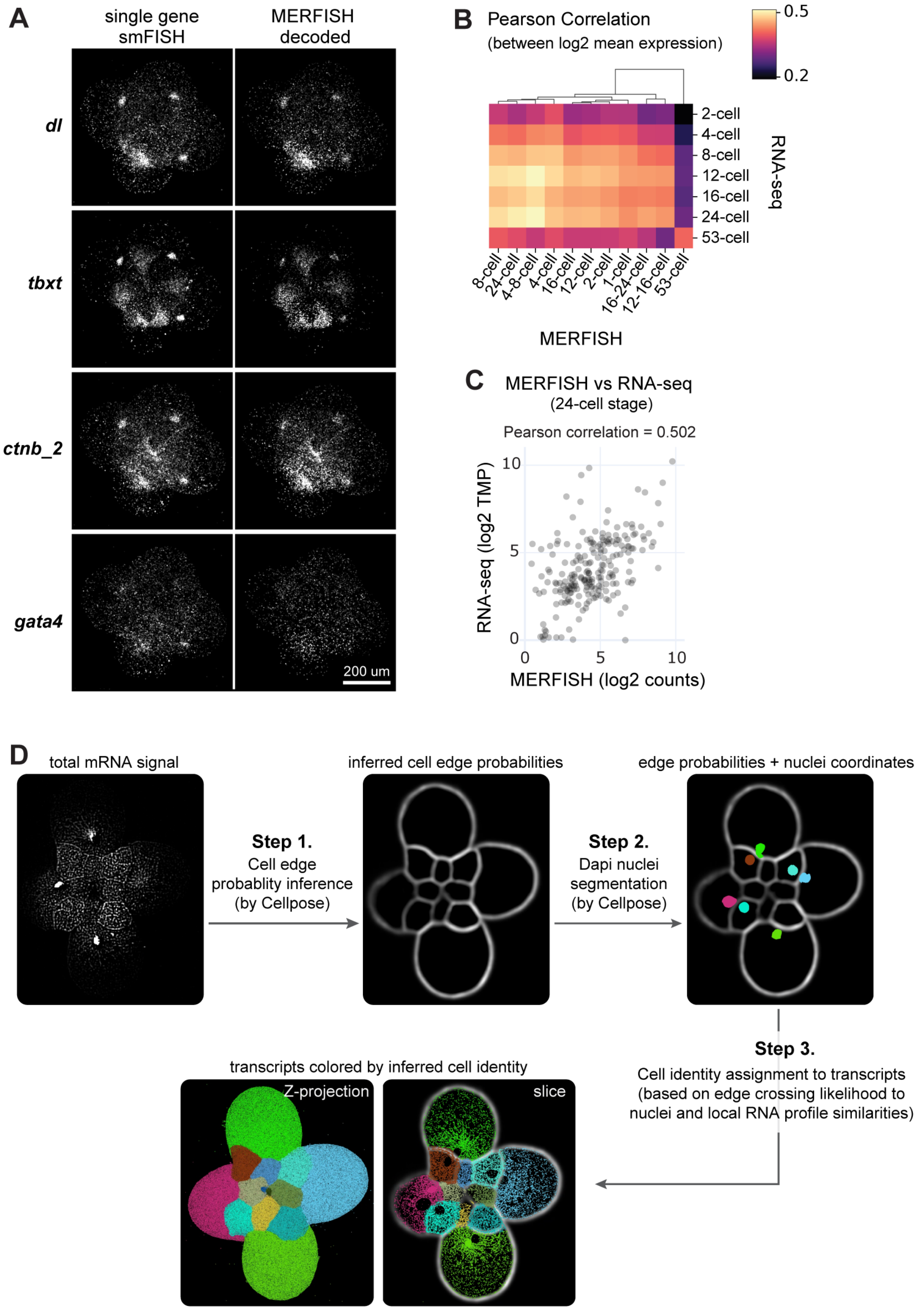


**Fig. S1. Quality check and cell segmentation of 3D MERFISH data in *C. atrasolea* embryos.** (**A**) Decoded MERFISH recapitulates single-gene smFISH results. The embryo shown is at the 24-cells stage. (**B**) Gene expression levels by MERFISH and bulk RNA-seq showed high correlations when measured at similar stages. Early cleavage stage embryos up to the 24-cell stage are transcriptomically similar and are distinct from late cleavage stage embryos (53-cells stage). (**C**) Scatter plot of gene expression levels measured by MERFISH versus bulk RNA-seq at the 24-cells stage. (**D**) Schematic of the cell segmentation pipeline used for assigning cell identities to MERFISH detected transcripts. A 12-cell stage embryo is shown in the example.

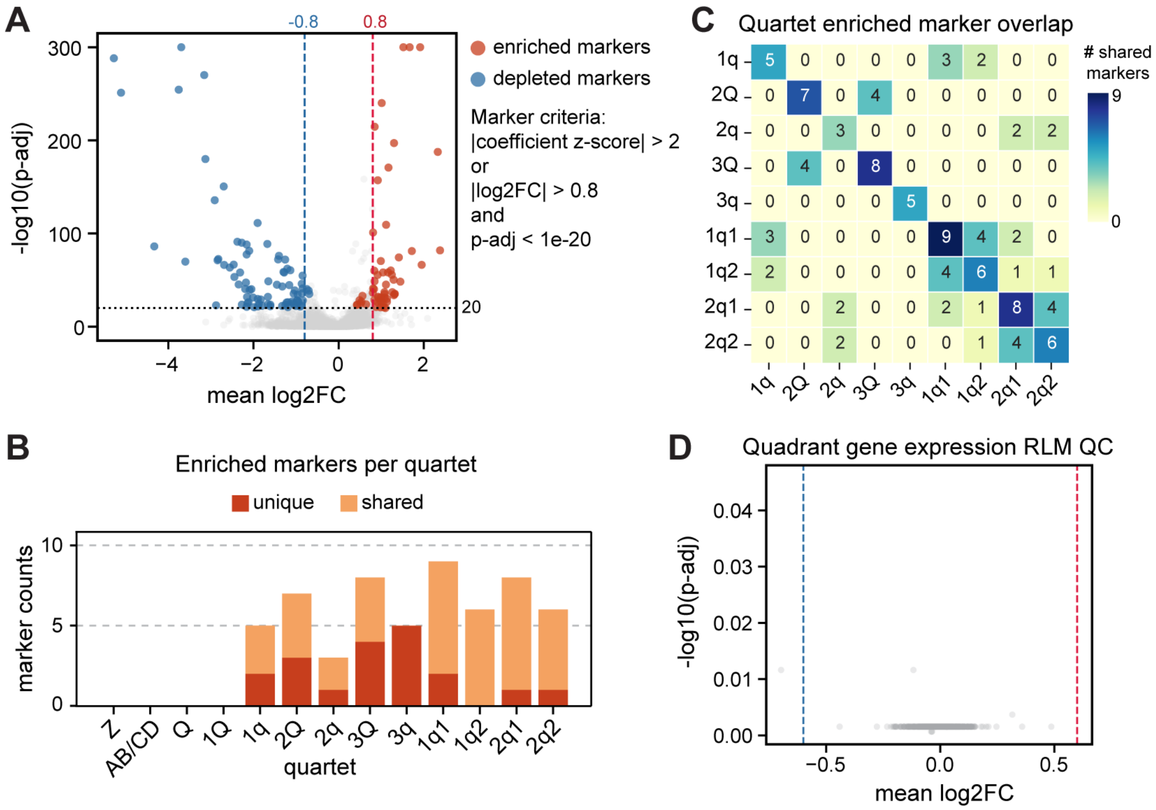


**Fig. S2. Lineage marker quantification.** (**A**) Volcano plot for quartet marker test. Each point represents a gene in a quartet. P-values are adjusted by Benjamini-Hochberg FDR correction. log2 fold change compares a gene's expression in the quartet to its mean expression across all other quartets. Genes that passed selection criteria for markers are highlighted in red (enriched) or blue (depleted). In addition to log2 fold change and adjusted p-value, a robust z-score was calculated per gene per lineage from the fitted RLM coefficients. This score serves as an alternative to fold change for measuring the extent of genes' differential expression. Genes passed either the score or fold change thresholds are considered markers if they also pass the p-value threshold. (**B**) Number of enriched marker genes (red genes in **A**) per quartet. (**C**) Number of shared enriched markers between quartet pairs. (**D**) Similar volcano plot as in **A** but for quadrant marker test. No significant markers were called due to low p-values and fold changes.

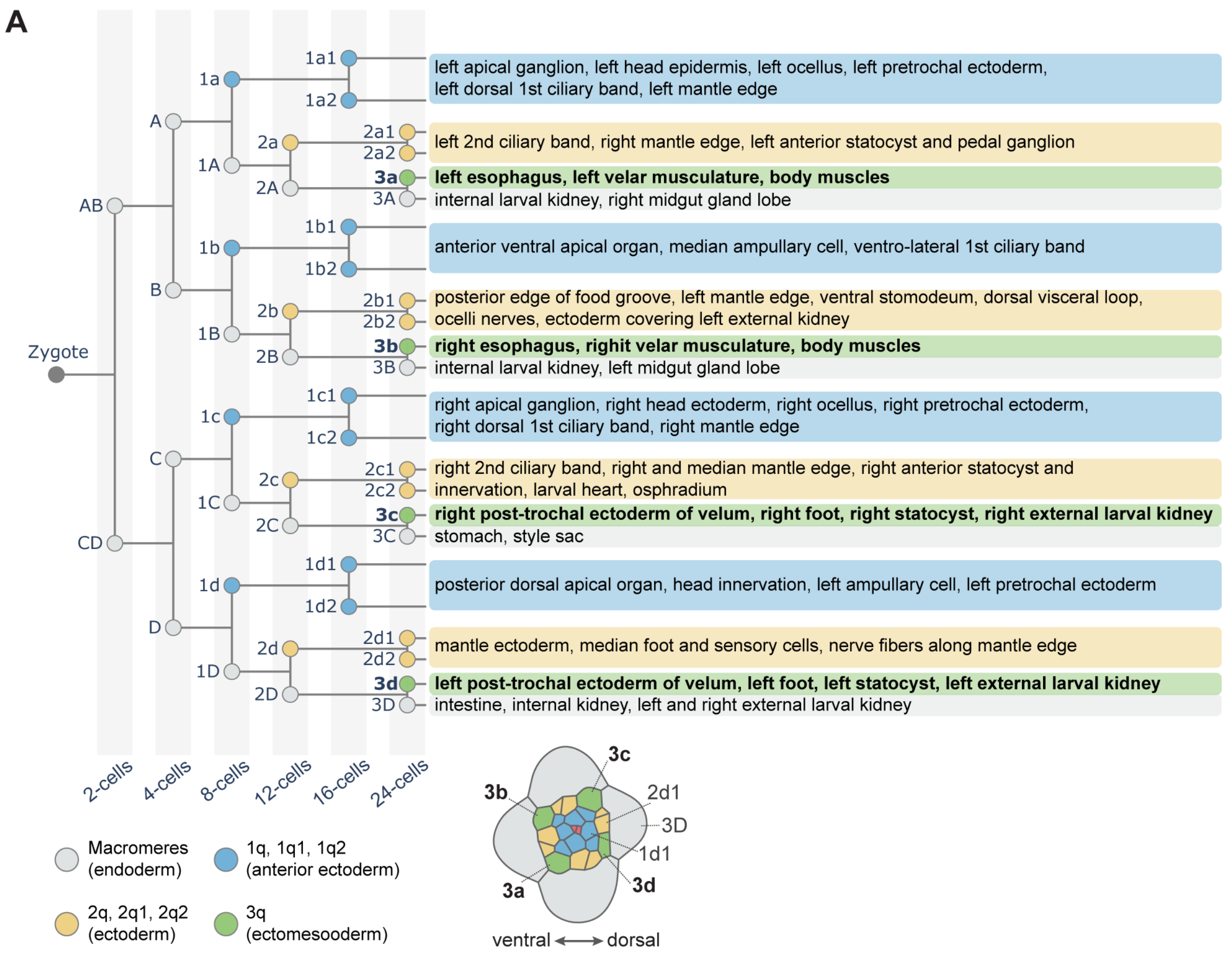


**Fig. S3. Cell lineage fatemap showing the larval fates of embryonic cells up to the 24-cells stage.** Shown is the lineage fatemap for *Crepidula fornicata*, a closely related sister species of *Crepidula atrasolea*. The lineage fatemap is based on ref. *31* and *36*. An illustration of a 24-cells stage embryo is shown at the bottom with cells colored by quartet groups. Annotations for the 3q cells (muscle progenitors) are bolded in the lineage fatemap and the embryo illustration.

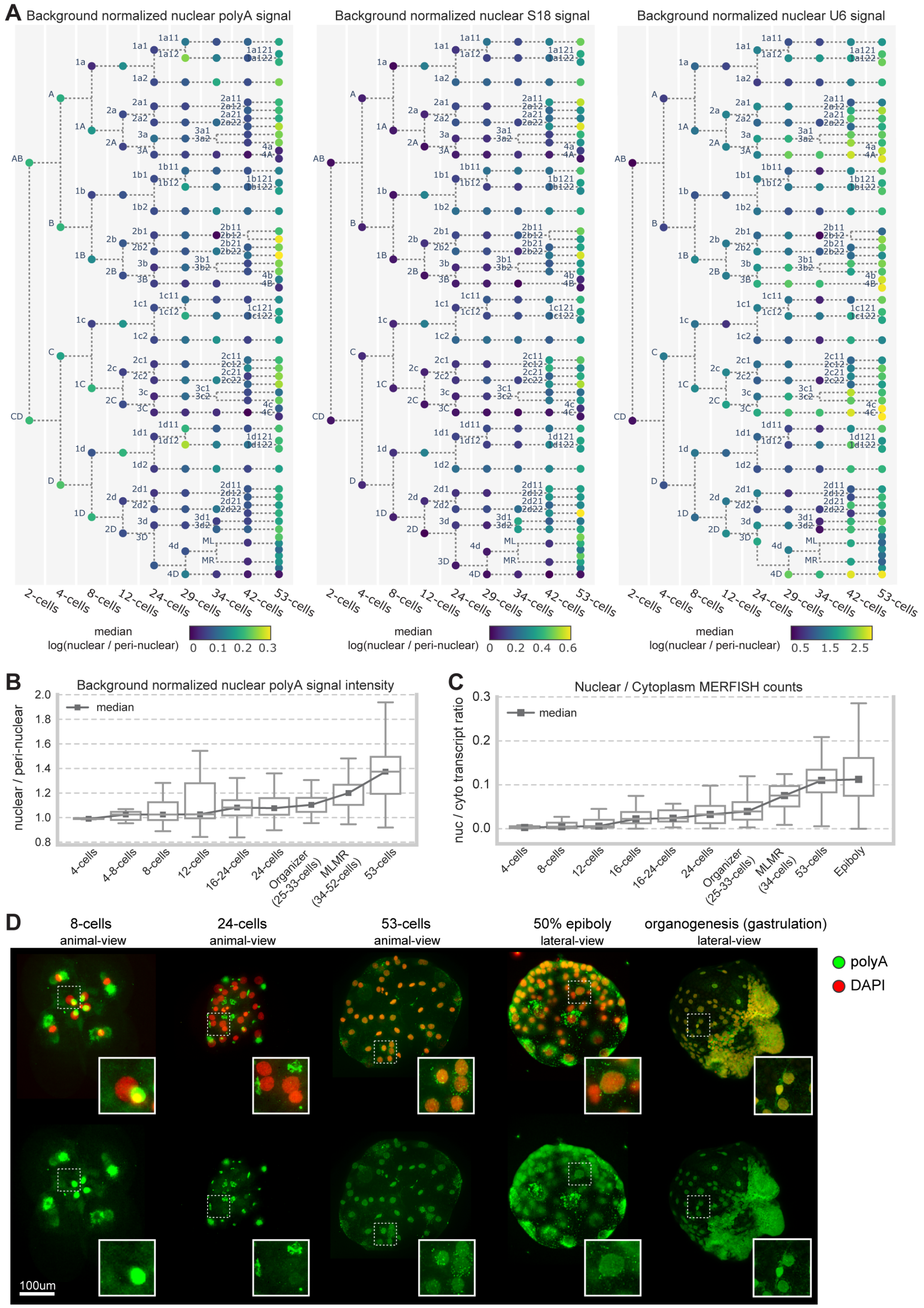


**Fig. S4. Nuclear RNA content is low until after the 24-cell stage.** (**A**) Median log background normalized nuclear signal from polyA FISH (left), S18 rRNA FISH (middle), and spliceosome component U6 FISH (right). (**B**) Quantification of normalized polyA FISH nuclear signal by stage. (**C**) Boxplot of nuclear transcript ratio from MERFISH data on the 200 genes assayed. (**D**) Nuclear mRNA content visualized by polyA FISH in embryos across early embryogenesis. Insets show representative interphase nuclei.

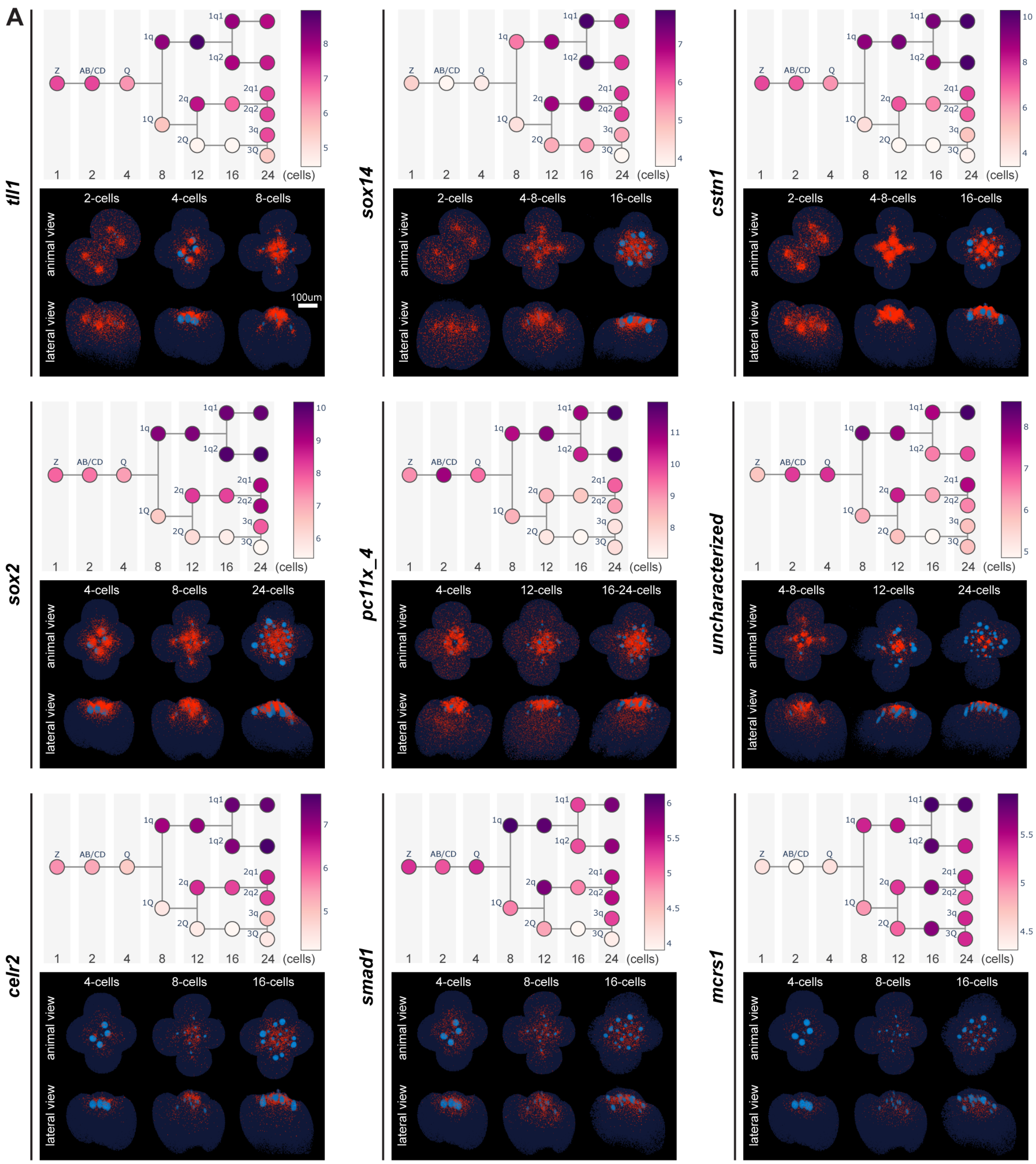


Fig. S5 (continued on next page).

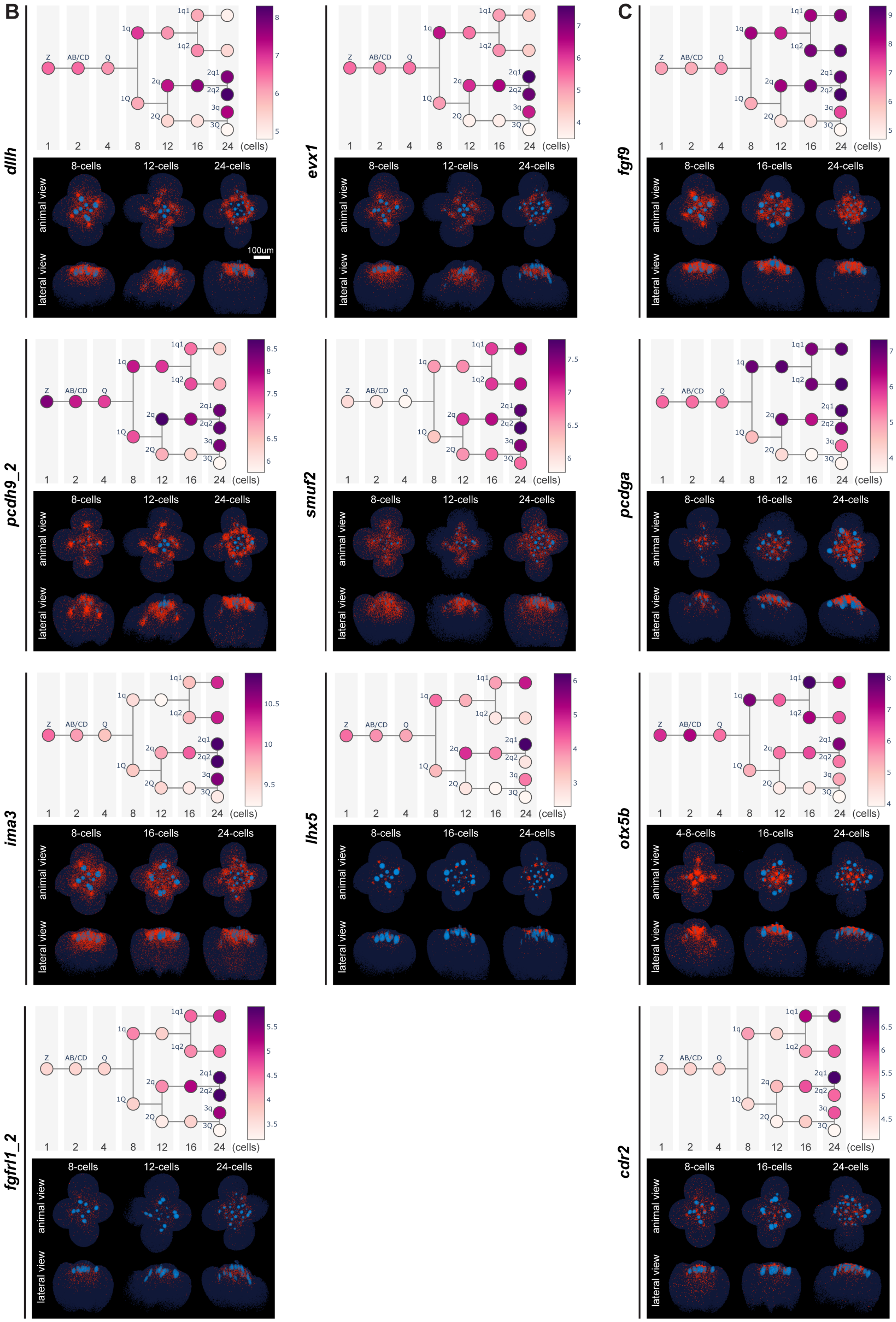


*Fig. S5 (continued on next page).*

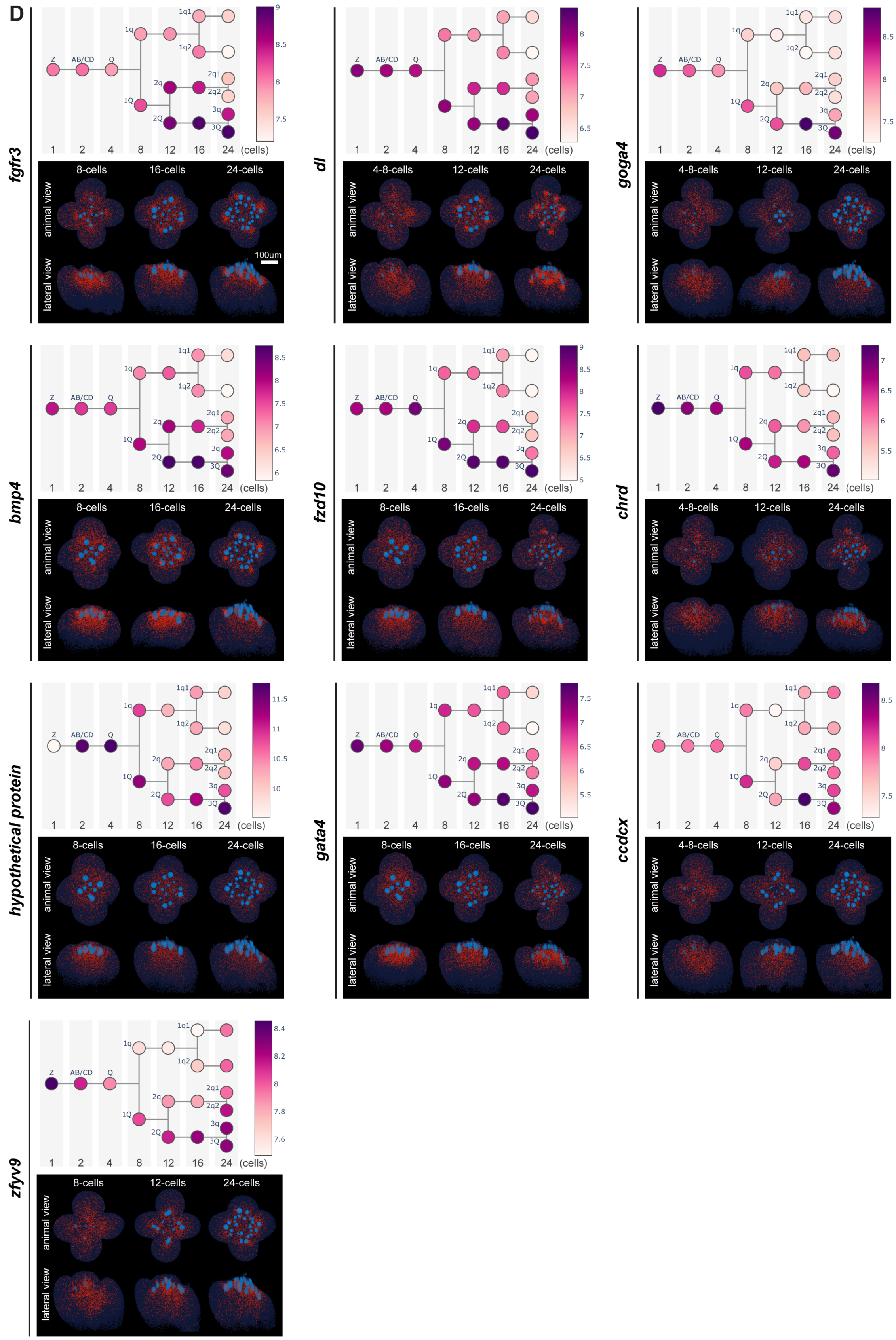


*Fig. S5 (continued on next page).*

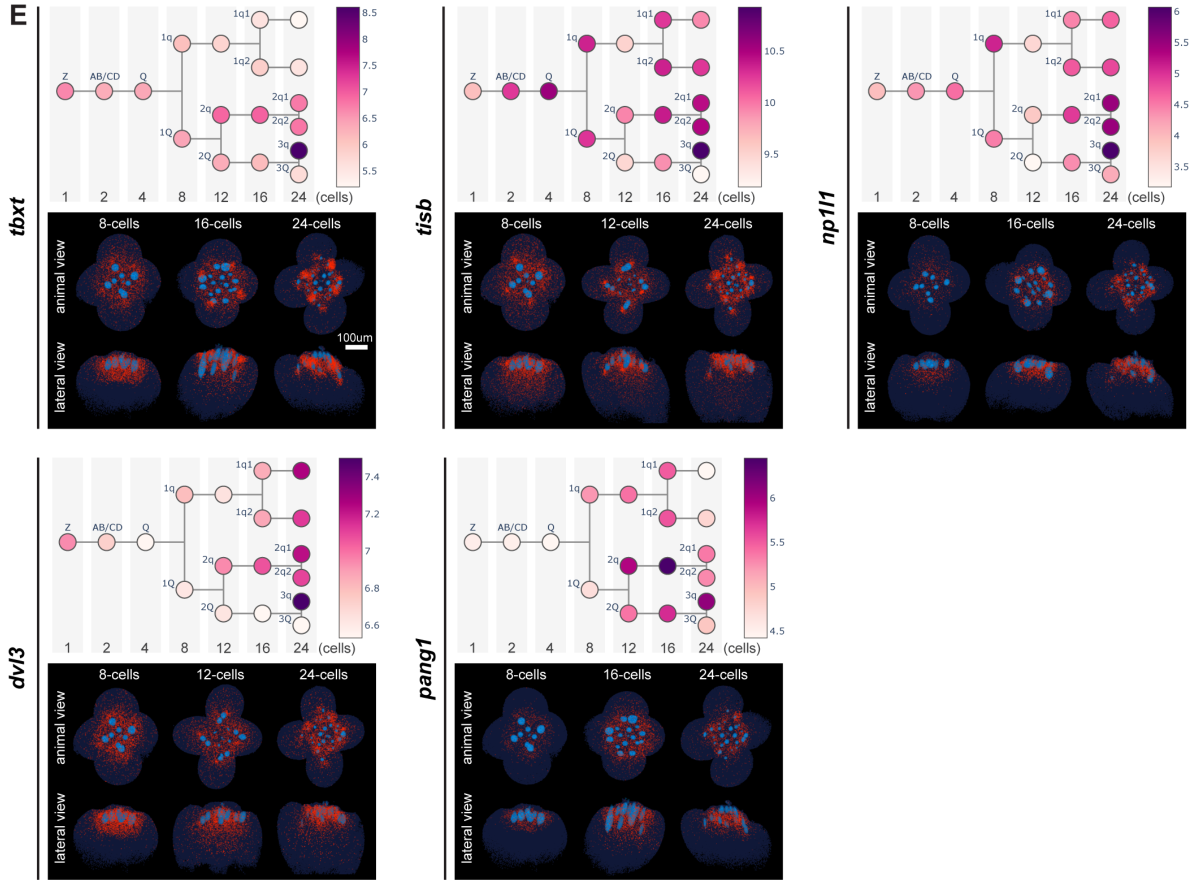


**Fig. S5. Quartet marker gene expression.** (**A**) Markers of the 1q lineages. (**B**) Markers of the 2q lineages. (**C**) Genes that mark both 1q and 2q lineages. (**D**) Markers of the macromeres. (**E**) Markers of 3q. For each gene, upper panel shows the log2 normalized counts overlaid on the cell lineage tree; lower panel shows smFISH style plots for selected embryos with marker gene transcripts in red, nuclei in bright blue, and overall embryo in dark blue. All embryos are oriented with the D quadrant to the right.

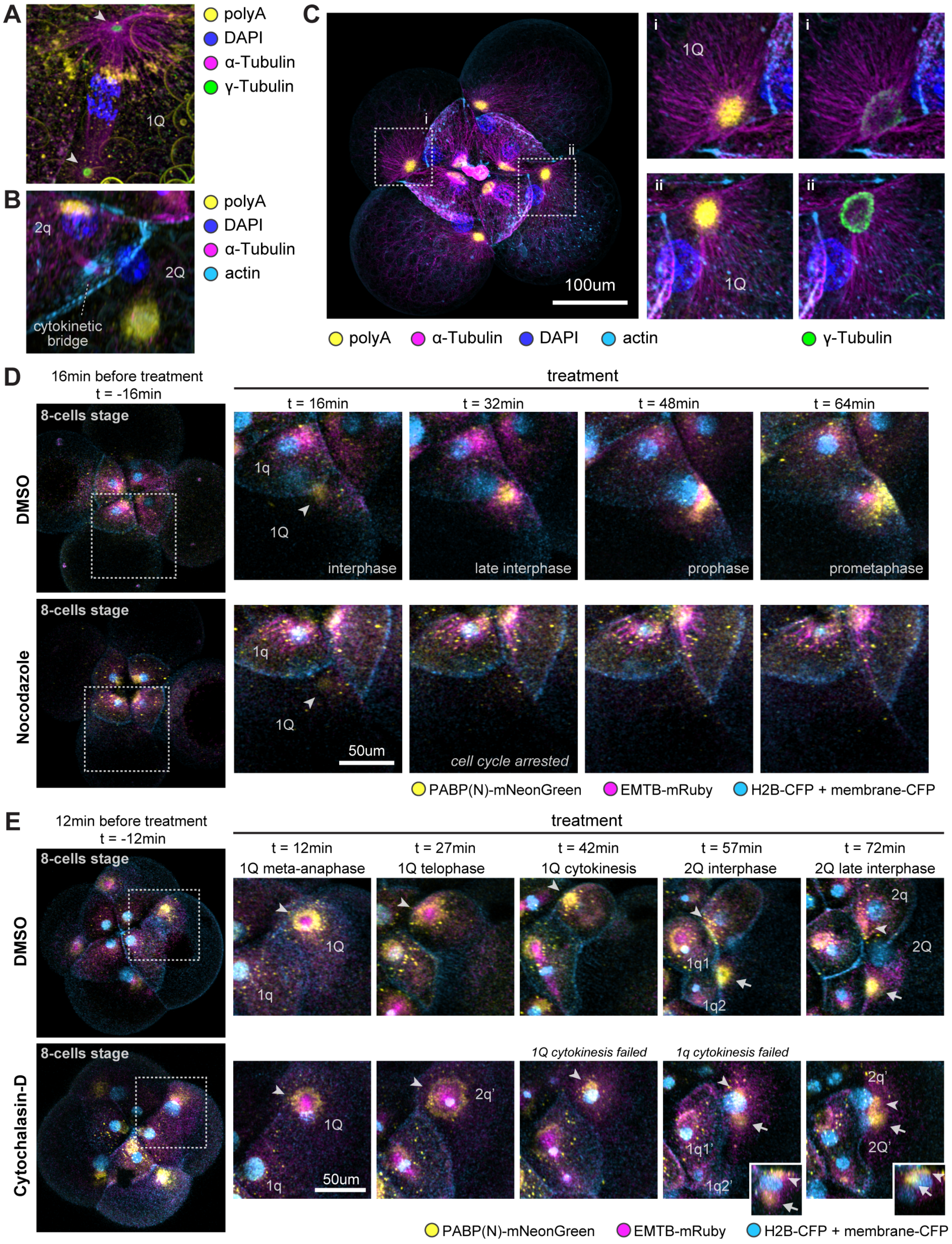


**Fig. S6. Total mRNAs are associated with microtubule structures.** (**A-C**) PolyA-FISH-coupled immunostaining. (**A**) mRNA aggregation (polyA) is colocalized with astral microtubules at 1Q anaphase. Arrowheads point to the centrosomes (marked by γ-tubulin). (**B**) mRNAs re-accumulate into primary aggregates upon completion of cell division. (**C**) The mRNA aggregate domain at late interphase is surrounded by γ-tubulin, suggesting that it is a MTOC. (**D**) Live-imaging of pharmacological inhibition of microtubule polymerization. Dashed boxes in pre-treatment images indicate regions shown in the snapshots during treatment. Animal projections are shown. Arrowheads point to mRNA aggregates formed in 1Q during interphase. Nocodazole treatment arrested the cell cycle and inhibited mRNA accumulation at the aggregate. (**E**) Live-imaging of pharmacological inhibition of actin polymerization. Dashed boxes in pre-treatment images indicate regions shown in the snapshots during treatment. Cytochalasin-D treatment abolished cytokinesis. 2q' and 2Q' labels the 2q and 2Q nuclei in the cell that failed to undergo cytokinesis. Small insets in the cytochalasin-D treatment are lateral projections of the 2q'/2Q' cell, with the animal side on top. Arrowheads point to the mRNA aggregate formed in 1Q and inherited by 2q (2q'). Arrows point to the mRNA aggregate formed in 2Q (2Q').

**Table S1. Genes assayed by MERFISH.** For each gene, the table includes its transcript ID, best blast hit, and its unique gene symbol used in the paper. (separate .csv file)

**Table S2. Marker genes of cell quartets.** Marker genes were identified by fitting a robust linear model (materials and methods). Each row records a marker gene, a cell group it marks, its log fold change, p-value, and an expression score. The score was calculated based on the fitted coefficient of a gene as a metric for the enrichment of the gene in a cell group. Some genes are listed in multiple rows, each corresponding to a different cell group they were identified as a marker for. (separate .csv file)

**Data S1. Probe sequences used for MERFISH in FASTA format.** This file contains the complete sequences of all probes ordered as an oligo pool. Each entry header contains the gene symbol, transcript ID, probe ID, and adaptor ID. Each probe contains primer sequences for probe amplification, adaptor sequences for readout probe binding, and primary probe sequence that hybridizes to the target gene (Materials and methods). (separate file)

**Movie S1. Live imaging of total mRNAs at early cleavage stages.** Shown is a representative movie from 5 imaged embryos across similar developmental stages. Yellow shows total mRNA visualized with PABP(N)-mNeon-green. Cyan shows both nuclei marked by H2B-CFP and cell membrane marked by LifeAct-CFP. Magenta shows microtubules marked by EMTB-TDmRuby. (separate file)

**Movie S2. Live imaging of total mRNAs at later cleavage stages.** Shown is a representative movie from 3 imaged embryos across similar developmental stages. Yellow shows total mRNA visualized with PABP(N)-mNeon-green. Cyan shows cell membrane marked by LifeAct-CFP. Red shows nuclei marked by H2B-RFP. (separate file)
